# Two-stage synaptic plasticity enables memory consolidation during neuronal burst firing regimes

**DOI:** 10.1101/2025.04.12.648539

**Authors:** Kathleen Jacquerie, Danil Tyulmankov, Pierre Sacré, Guillaume Drion

**Author notes:** **Corresponding authors** Kathleen Jacquerie. **Declaration of interests** The authors declare no competing interests. **Declaration of generative AI and AI-assisted technologies** ChatGPT was used to correct for spelling. **Author contributions** Conceptualization, K.J., G.D.; Methodology, K.J., D.T., G.D.; Software, K.J.; Writing – Original Draft, K.J.; Writing – Review and Editing, K.J., G.D., P.S., D.T.; Visualization, K.J.; Supervision, G.D., P.S., D.T.; Funding Acquisition, K.J., G.D., P.S.

## Abstract

Neural circuits routinely alternate between input-driven tonic activity and collective burst firing. In the presence of Hebbian plasticity, bursts generate a robust attractor in weight space, creating a built-in drift that can be repurposed into a stabilizing trace of prior learning. We show that this phenomenon can be harnessed for memory consolidation through the introduction of a two-stage synaptic rule. The effective synaptic weight is defined as the product of a primary weight—updated by a Hebbian rule during both tonic and burst periods—and a secondary weight that updates in proportion with a coupling gain to the negative time-derivative of the primary weight. In a MNIST-like task, alternating tonic and burst epochs preserves earlier patterns, improves generalization to unseen inputs, and resists interference and noise, whereas replacing burst by quiescence or additional tonic epochs does not. Parameter sweeps reveal that coupling gain and the initial synaptic weights control whether bursts consolidate (“up-selection”) or prune (“down-selection”) synapses. Pairing the rule with alternative primary plasticity models yields distinct treatments of overlapping inputs, enabling either integration or separation. Studying switches in firing activity with a two-stage synaptic plasticity provides a plausible route to consolidation in biological and neuromorphic networks.

**Significance Statement:** Neural circuits alternate between tonic spiking and burst firing, yet most models of synaptic plasticity are limited to a single firing regime. We introduce a two—stage synaptic rule in which a primary weight encodes activity during both states, while a secondary weight—engaged only during bursts— stabilizes learning from tonic periods. In conductance-based networks and a pattern recognition task, this rule preserves memories, improves generalization, and resists interference, whereas quiescence or extended tonic activity do not. The model further shows that bursts can consolidate or prune synapses depending on coupling gain and initial conditions. These findings identify a plausible, biologically motivated mechanism for how activity state transitions shape memory consolidation.

**Graphical abstract:** 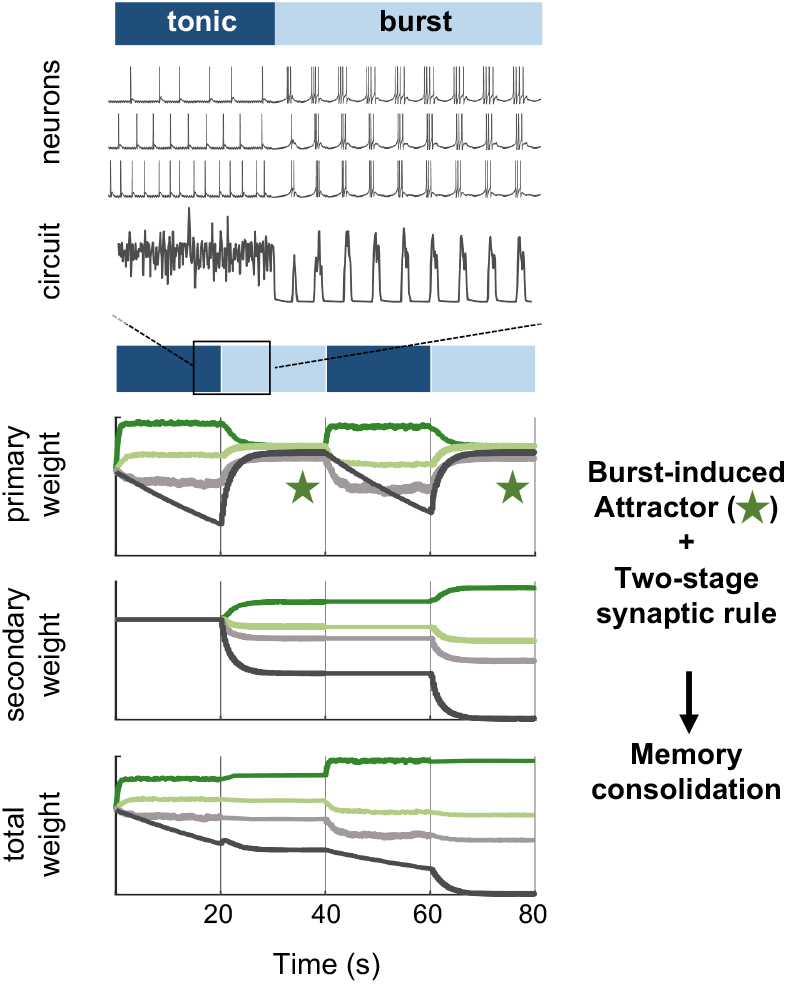

## 1 Introduction

Neural circuits naturally fluctuate between asynchronous tonic firing and collective bursting, depending on behavioral context and neuromodulatory state (McGinley et al., 2015). In tonic firing, neurons emit relatively isolated spikes that are primarily driven by ongoing external inputs. Burst firing consists of rapid clusters of spikes followed by periods of silence. Bursts can emerge endogenously from intrinsic neuronal and network properties, reflecting cycles of ion channel activation and inactivation, like T-type calcium channels (McGinley et al., 2015; Jacquerie and Drion, 2021). When many neurons burst synchronously, the resulting collective dynamics generate large-amplitude, low-frequency signal in local field potential (LFP). These transitions are widespread across brain regions—from thalamus to hippocampus—and shape how information is encoded and stabilized (McGinley et al., 2015; Sherman, 2001; Jacquerie et al., 2025).

Models of synaptic plasticity, however, treat each of these firing regimes in isolation, capturing synaptic changes in either tonic spiking or burst firing (Gjorgjieva et al., 2009; Costa et al., 2015; Clopath, 2019), but rarely how learning unfolds when networks switch between them. Tonic firing encodes external inputs and drives synapses toward a broad set of convergence values. Meanwhile, as shown in our previous work (Jacquerie et al., 2025), when a network enters collective burst firing, synaptic weights are pulled into a narrow region of weight space—a burst-induced attractor—arising due to induced similarity in either calcium dynamics or spike-time correlations across synapses throughout a network. This burst-induced attractor reorganises the weight space and provide a stabilizing trace of prior tonic learning. The existence of this phenomenon raises the question: how can a net-work preserve or consolidate information acquired during tonic firing while still benefiting from the reshaping of the weight space into an attractor during burst firing?

Taking inspiration from the idea of traces in plasticity (Fusi et al., 2005; Gerstner et al., 2018), we note that burst firing itself provides a built-in trace by reorganizing synaptic weights. We exploit this property by introducing a biologically plausible, analytically tractable, and computationally lightweight *two-stage plasticity rule*. In this framework, the effective synaptic strength is expressed as the product of two variables, which we term the primary and secondary weights. The *primary weight* follows a classic calcium-based plasticity rule (Graupner and Brunel, 2012), active during both tonic and burst states. The *secondary weight* changes only during bursts, according to a novel rule we propose in this work—negatively proportional to the rate of change of the primary weight, controlled through a *coupling gain*.This coupling transfers information from the primary to the secondary weight as the primary converges to the burst-induced attractor. Once the primary weight stabilizes at its attractor, the secondary weight also remains stable. This formulation offers a plausible mechanism for modeling the biological signaling cascades that operate in the postsynaptic compartment. Calcium fluctuations drive rapid changes, for example, by regulating postsynaptic receptor activation or AMPA receptor insertion (Lamprecht and LeDoux, 2004). In turn, these changes trigger downstream signaling pathways inside the synapse (Citri and Malenka, 2008).

We evaluate this plasticity rule in biophysical conductance-based neural networks: first, in a small circuit to demonstrate its dynamics and enable tractable analysis, then in a simple pattern recognition task to show a proof-of-concept of its practical utility. Alternating tonic and burst states with the two-stage rule enables memory consolidation: learned patterns remain stable, and even generalize to novel inputs. In ablation experiments—with no secondary plasticity, extended input-driven tonic firing, or quiescent intervals of sparse tonic firing instead of bursts—synapses fail to consolidate and quickly forget previously stored information. We further show that the coupling gain and initial conditions of the synaptic weight can determine whether bursts lead to consolidation (up-selection) or pruning (down-selection), providing a flexible mechanism for regulating stability and plasticity. Finally, we test how modifications to the primary synaptic rule shape different strategies for learning overlapping patterns.

Overall, this work introduces a minimal biologically grounded synaptic rule that uses network state transitions for memory consolidation. By combining a primary calcium-based plasticity with a secondary plasticity rule, our model inherits both the flexibility of tonic firing as together with the robustness of burst-driven stabilization.

## 2 Results

### 2.1 Modeling robust transitions between tonic and burst firing in a biophysical network model

To model a neural circuit that can switch between tonic and burst firing regimes, we use a conductance-based spiking network introduced in our previous work (Jacquerie et al., 2025). The architecture consists of *N* presynaptic excitatory neurons projecting to *M* postsynaptic excitatory neurons, all receiving inhibition from a single pacemaking inhibitory neuron. The inhibitory neuron regulates the network state, mimicking the role of neuromodulators such as acetylcholine, dopamine, serotonin, or histamine—known to control brain state transitions (Hill and Tononi, 2005; Bazhenov et al., 2002; Olcese et al., 2010; Jacquerie and Drion, 2021; Drion et al., 2018). When depolarized, it induces *input-driven tonic firing* in the excitatory neurons, which spike independently of each other in response to their external input currents. When hyperpolarized, it induces *collective burst firing*, in which the excitatory neurons generate synchronized clusters of spikes followed by a silent period, driven by intrinsic conductances such as T-type calcium channels. At the population level, these two regimes produce distinct local field potential (LFP) signatures: fast, low-amplitude oscillations during tonic activity and slow, high-amplitude oscillations during burst firing (Figure 1A–B). This switching mechanism provides a biologically plausible framework for studying how synaptic plasticity operates across alternating neural activity states.

**Figure 1:**
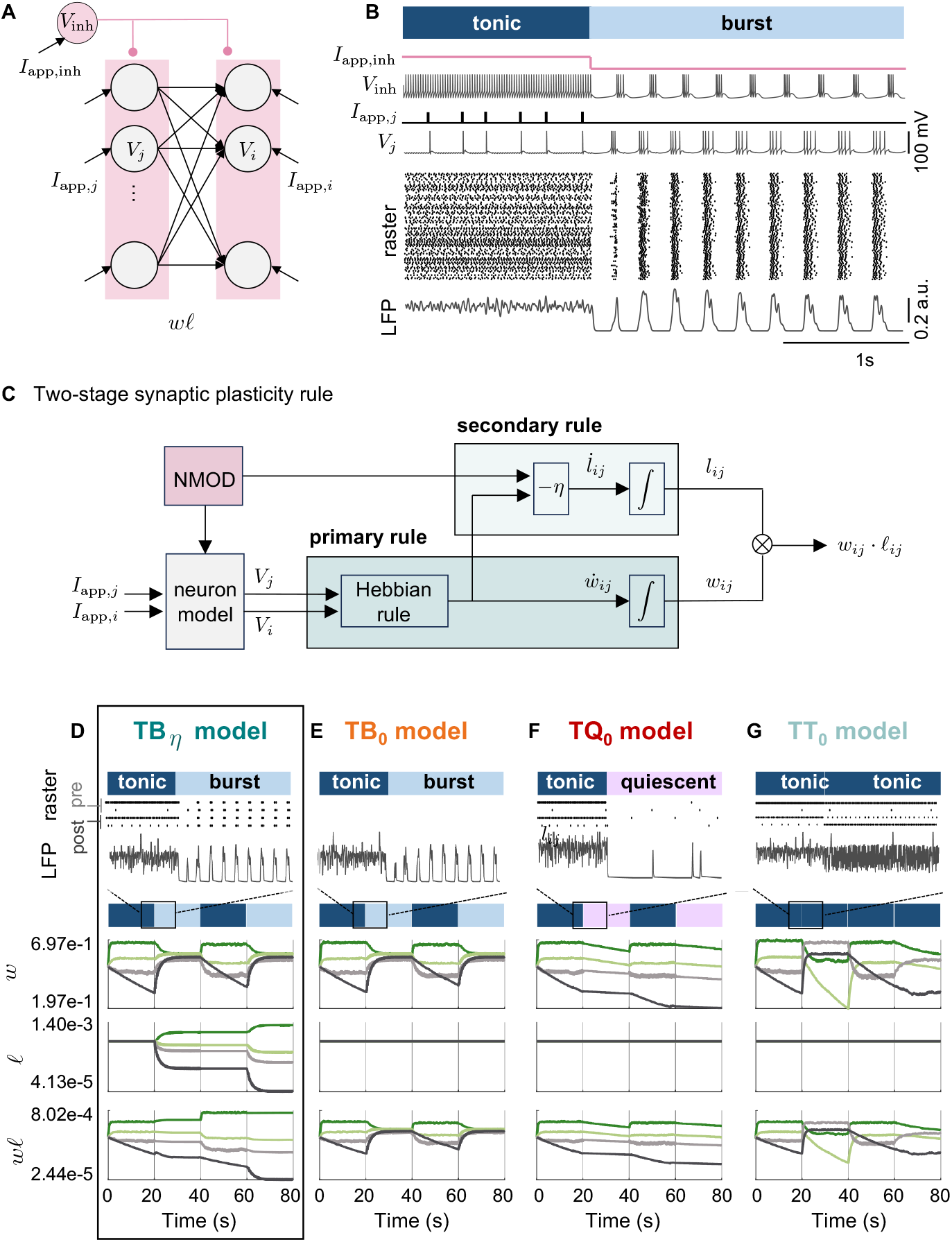
A biophysical network model of tonic–burst transitions with two-stage synaptic plasticity. **A**. Network architecture: one inhibitory neuron (*V*_inh_) projects to all excitatory neurons, which are connected feedforward from *N* presynaptic neurons (*V*_*j*_) to *M* postsynaptic neurons (*V*_*i*_). An external current to the inhibitory neuron (*I*_inh_) controls the network state; each excitatory neuron receives an independent external current mimicking input (from (Jacquerie et al., 2025)). **B**. Hyperpolarizing *I*_inh_ switches the network from input-driven tonic firing to collective burst firing. Shown are applied currents, membrane potentials, raster plots, and LFP traces (A–B adapted from (Jacquerie et al., 2025)). **C**. Two-stage synaptic plasticity illustrated in the state diagram. Primary plasticity (*w*_*ij*_) operates during both tonic and burst states, following a Hebbian rule driven by pre- and postsynaptic activity, we mainly use the calcium-based model (Graupner and Brunel, 2012; Graupner et al., 2016). Secondary plasticity (*𝓁*_*ij*_) is active only during burst firing and changes proportionally and oppositely to the primary weight derivative, scaled by coupling gain *η*. The total weight is the product between the primary and the secondary weights. Neuromodulation (NMOD) can switch the firing pattern of the neurons, turn on-and-off or tune the coupling gain (*η*). **D–G**. Evolution of primary (*w*), secondary (*𝓁*), and total (*w𝓁*) weights in a two-by-two neuron circuit with varying correlation levels. **D**. Tonic-burst transition with with *η* = 1*/*500: secondary weights store tonic-encoded learning; total weights consolidate during bursts. **E**. Tonic-burst transition with *η* = 0: only primary plasticity acts; burst firing drives weights toward the burst-induced attractor. **F**. Tonic-quiescent transition (*η* = 0): quiescence replaces burst firing; primary weights decay slowly. **G**. Tonic-tonic transition (*η* = 0): input-driven tonic replaces burst firing; primary weights encode new memories.

### 2.2 A novel coupled two-stage synaptic plasticity rule

The synapses projecting from the presynaptic to the postsynaptic excitatory neurons undergo plasticity. In (Jacquerie et al., 2025), we demonstrate that using either a traditional calcium-based rule or an STDP rule during collective burst firing leads to the presence of a burst-induced attractor, driving all the synaptic weights towards the same narrow region in the weight space. In this work, to protect memories from being corrupted during ongoing activity and storage of new memories, we take inspiration from (Fusi et al., 2005; Benna and Fusi, 2016) to construct models of synaptic plasticity which couple different stages to mimic the dynamic biochemical systems ubiquitous in biological systems.

We build a two-stage synaptic model where the total synaptic weight is the product of a primary weight (*w*_*ij*_) and a secondary weight (*𝓁*_*ij*_). The primary weight follows a Hebbian plasticity rule active during tonic and burst firing (Figure 1C). The rate of change of the primary weight is a function *f* of the presynaptic neuron *j* and the postsynaptic neuron *i* such as:

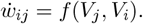

We use a calcium-based rule as an example (Graupner and Brunel, 2012; Graupner et al., 2016).

The secondary weight is modeled such as the rate of change of the secondary weight is proportional to the negative rate of change of the primary weight:

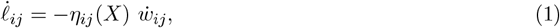

where the parameter *η*_*ij*_(*X*) ≥ 0 represents the coupling gain between presynaptic neuron *j* and postsynaptic neuron *i*. In the most general case, it can be a function of neuromodulatory or other circuit parameters, denoted as the vector *X*, allowing its value to differ across circuit regimes. In our model, we consider *X* = *I*_app,inh_, corresponding to the neuromodulatory signal controlling the transition between tonic and burst firing, and a uniform gain across all synapses in the circuit:

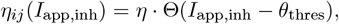

where Θ(·) is the Heaviside step function, such that the secondary rule is engaged only during bursts (*i*.*e*., when *I*_app,inh_ is above a threshold *θ*_thres_) and 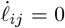 during tonic states. See Methods subsub-section 4.2.6 for more details.

For simplicity, we assume that the coupling gain is uniform across the network and denote this global value as *η*. Increasing *η* produces stronger modifications of the secondary weight, reflecting a tighter coupling between cascade mechanisms (Figure 1C). The effect of varying *η* is analyzed in subsection 2.6, while the possibility of modulation at the level of individual synapses or subnetworks is considered in the Discussion.

The model makes two hypotheses. First, there exists a negative proportional relationship between the rates of change of the primary and secondary weights. This formulation captures the idea that learning acquired during tonic firing is reflected in the primary weight, while burst firing triggers a complementary process that transfers this information into the secondary weight. The negative sign in the secondary plasticity equation ensures that, as the primary weight converges toward the attractor, its changes are mirrored in the secondary weight. This mechanism allows the total synaptic weight to preserve previously encoded information. Second, this secondary mechanism is engaged specifically during burst firing. This assumption is motivated by the observation that transitions into bursting are often accompanied by neuromodulatory signals, which can in turn activate biochemical cascades that support longer-lasting changes in synaptic efficacy.

### 2.3 Synaptic weight trajectory across tonic, burst, quiescent transitions

We first examine how the proposed two-stage plasticity rule influences the memory storage in a simple circuit. To do so, we simulate a minimal circuit with two presynaptic and two postsynaptic neurons. The presynaptic neurons fire at distinct tonic frequencies, producing heterogeneous synaptic trajectories during tonic activity (see Methods 4.2.7 for details).

In the tonic–burst model with coupling (TB_*η*_ model, Figure 1D, *η* = 1*/*500), the network switches from tonic to burst firing while secondary plasticity is active. During tonic firing, the primary weights *w* encode the different inputs. When the network enters bursting, the primary weights converge toward the burst-induced attractor. Because the secondary weight *𝓁* tracks the negative derivative of *w*, the convergence of the primary weights towards the attractor lead to a transfer of information into the secondary weights *𝓁*. Once *w* reaches the attractor, *𝓁* stabilizes as well, so that the total effective weight (*w𝓁*) preserves the memories acquired during tonic firing.

To assess the role of the two-stage plasticity, we disable the coupling by setting *η* = 0, leaving only the primary Hebbian mechanism (TB_0_ model, Figure 1E). In this case, tonic firing increases the primary weights, but the subsequent burst drives all *w* values toward the attractor. Without secondary plasticity, this convergence erases previously encoded information, leaving only transient traces. Biologically, this resembles blocking cascade signaling pathways required for long-lasting plasticity (Frey et al., 1993; Krug et al., 1984).

We next ask what happens if burst firing is replaced by a quiescent regime. This condition reflects a common modeling practice, where “offline” states are approximated by inactivity (Fauth and Van Rossum, 2019). In the tonic–quiescent model (TQ_0_ model, Figure 1F, *η* = 0), synaptic weights change during tonic firing, but the subsequent quiescent phase provides no mechanism for stabilization. As a result, the weights decay slowly and retain only short-lived information.

Finally, we test the effect of replacing burst episodes with additional tonic learning. In the tonic–tonic model (TT_0_, Figure 1G, *η* = 0), each burst period is substituted by another tonic phase in which new inputs are presented. Instead of consolidating prior information, these additional tonic episodes overwrite previously encoded patterns, preventing memory stabilization.

Together, these experiments aim to study how the synaptic weights evolve when the secondary rule is ablated or when the burst firing is replaced by either sparse tonic or additional tonic firing encoding new digits. These comparisons highlight that switches between tonic and burst firing with two-stage plasticity is uniquely suited to preserve and stabilize memories: bursting alone triggers the burst-induced attractor, quiescence allows only transient storage, and continuous tonic firing induces interference. In contrast, when including the two-stage rule, burst episodes actively consolidate tonic-encoded information.

### 2.4 The two-stage synaptic plasticity promotes pattern consolidation during burst firing

We next test whether the proposed two-stage synapse model supports memory consolidation in a pattern recognition task. The network is trained on the MNIST dataset of handwritten digits (LeCun et al., 1998), using downsampled 22 × 22 images for computational efficiency. Each of the *N* = 484 presynaptic neurons encodes the intensity of a single pixel by its firing rate, while *M* = 10 postsynaptic neurons represent the ten digit classes in a “one-hot” encoding.

Training consists of 21 sequences (Figure 2A, Roman numerals I–XXI), with each sequence composes of 10 blocks. For sequences I through XX, in block *i* (for *i* = 0, 1, …, 9), a randomly sampled image of digit *i* is presented to the network during the input-driven tonic firing (Figure 2A, dark blue, T), followed by either collective bursting (Figure 2A, light blue, B), quiescence (Figure 2A, pink, Q), or an additional tonic firing period (T’), as in the previous section. The final sequence XXI test robustness by presenting noise input (N). During tonic firing, synapses update according to the primary calcium-based plasticity rule (Graupner and Brunel, 2012). During burst firing, the secondary weight is upated according to the two-stage plasticity (TB_*eta*_ model). For more details about the implementation, see Methods 4.2.8.

**Figure 2:**
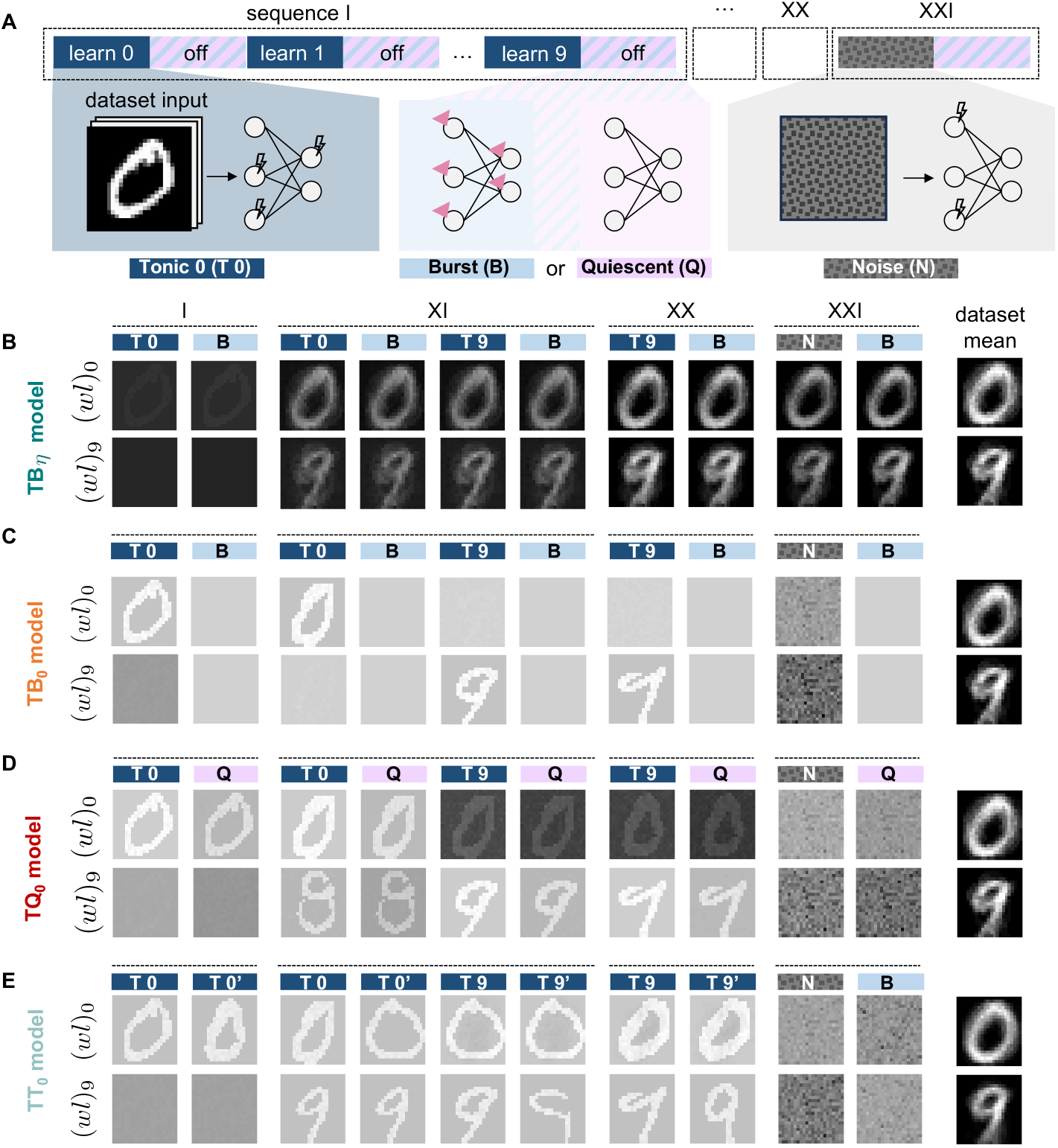
Memory consolidation through burst firing and two-stage plasticity. **A**. Schematic of the experimental protocol. The network sequentially encodes digits (0–9) during input-driven (shown by the lightning symbol) tonic firing periods (T) (interleaved with off period (hatched lavender, blue), which can be modeled either as burst firing (B) or quiescence (Q). It consists in 2° sequences (I-XX). In the final sequence, random noise is presented (N) on the last sequence (XXI). **B-E**. The weight matrices are shown for the output neurons associated with digits 0 and 9 at the transition from tonic to burst firing within one sequence (noted (*w𝓁*)_0_ and (*w𝓁*)_9_). The grey color scale is normalized between the minimum and maximum weight of the simulation. The last column is the mean of the 20 dataset images for the digits 0 and 9 presented during the 20 sequences. **B**. Tonic–burst with two-stage plasticity (TB_*η*_, *η* = 1*/*400): secondary weights preserve information learned during tonic firing, enabling the total weight (*w𝓁*) to retain and consolidate memories across sequences. **C**. Tonic–burst model (TB_0_, *η* = 0): During burst firing, primary weights converge to the burst-induced attractor, leading to loss of previously encoded patterns. **D**. Tonic–quiescent model (TQ_0_, *η* = 0): quiescence replaces bursts; weights decay gradually and memories are not consolidated. **E**. Tonic–tonic model (TT_0_, *η* = 0): off periods are replaced by additional tonic firing periods for sequences I to XX (T *i*’). The network only encodes the current digit, forgets previous learning, and is fragile to noise. The last column is the mean of the 40 dataset images presented during the 40 tonic firing states.

To visualize learning progress, we monitore the set of synaptic weights associated with each postsynaptic neuron—corresponding to a unique digit. The set of effective synaptic weights for digit *i* is defined as the element-wise product of primary and secondary weights afferent onto neuron *i*, in other words (*w𝓁*)_*i*_ = *w*_*i*:_ ⊙*𝓁*_*i*:_. For visualization, this set of weights is then reshaped into a 22 × 22 matrix to recover the pixel structure. Examples for digits 0 and 9 are shown in Figure 2B–E (all digits in Figure S1).

The results highlight the synergic role of bursting and the two-stage plasticity. The TB_*η*_ model (*η* = 1*/*400) preserves information: the secondary weight transferred learning from tonic periods into a stable component, enabling the total weight to retain clear digit representations across sequences (Figure 2C). This demonstrates memory consolidation without requiring replay of stored patterns. We also observe a phenomenon of generalization, where the network forms a combined representation of the different digit samples. This is evident in the weight matrices, which do not reflect only the most recent digit, but instead integrate features from the samples presented during earlier tonic firing states. To assess the role of the two-stage plasticity, we disable the coupling by setting *η* = 0, leaving only the primary Hebbian mechanism (TB_0_ model). Burst firing erases previously learned patterns by driving weights toward the burst-induced attractor (Figure 2B, shown by the grey pixel structure). In comparison, we replace burst firing by a quiescient periods, use in computational models to reproduce inactivity in a network. The TQ_0_ model only encodes the current digit, then leads to gradual weight decay and poor retention (Figure 2D). Additional learning can be seen as an appealing strategy for learning more information instead of alternating between tonic and burst firing. However, we observe that the TT_0_ model (tonic-only) overwrites previously stored digits with each new input, preventing consolidation and leaving the network fragile to noise (Figure 2E).

Together, these experiments show that consolidation emerges only when burst firing is combined with the two-stage rule.

### 2.5 The two-stage synaptic plasticity rule supports memory retention and generalization

To quantify the learning process, we compute the correlation between the weights encoding each digit and its template—the mean image of the corresponding digit class in the training dataset (the mean of the 20 training images for digits 0 and 9 is shown in the last column of Figure 2B–E). A high correlation indicates that the synaptic weight pattern closely resembles the template of that digit, meaning that the network retains the dataset mean of samples; a low or negative correlation indicates that the representation has been lost or overwritten. This measure provides a direct and interpretable proxy for memory retention between the different network states.

In the TB_*η*_ model, the correlation for each digit increases steadily as the network trains and remains high across sequences, approaching 1 even in the presence of noise (Figure 3A). Ablating the secondary rule by setting *η* to 0 during burst leads to a rising correlation during tonic firing that drops as the weights are pulled toward the burst-induced attractor during bursts (TB_0_ model, Figure 3A). Replacing burst firing with quiescent activity (TQ_0_ model) produces an initial increase in correlation upon digit presentation, but this quickly fades during subsequent states (Figure 3C). In the TT_0_ model, where off periods are replaced with additional tonic learning, the correlation rises when a digit is presented but then falls when the next digit is introduced (Figure 3D). This confirms that the network fails to maintain previously learned representations when bursting is absent.

**Figure 3:**
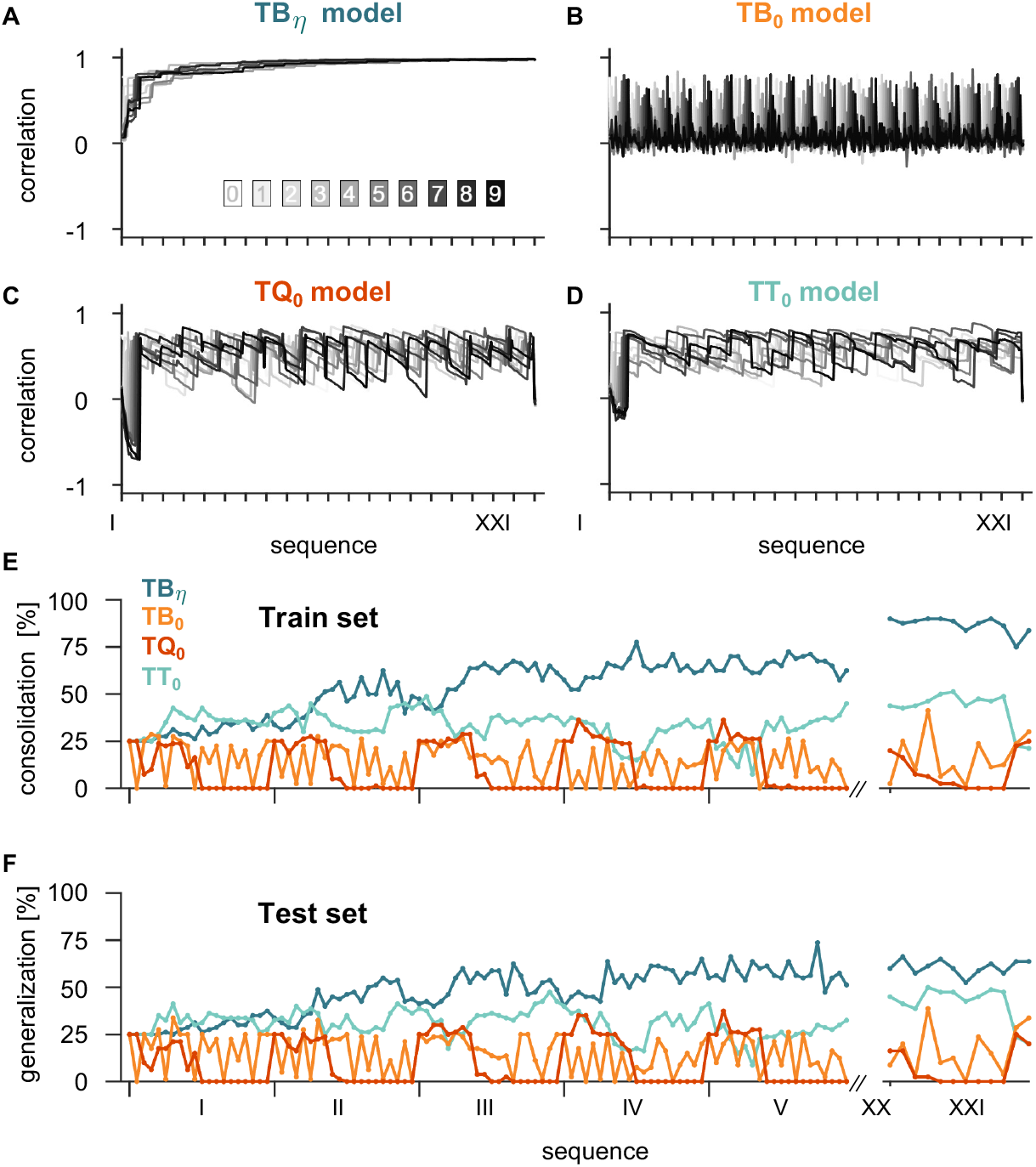
The two-stage synaptic plasticity rule supports memory retention and generalization. **A–D**. Correlation between the synaptic weight matrices and the average digit images from the training dataset across learning sequences (I–XXI). Each trace corresponds to one digit class (0–9), shown in grayscale from light (digit 0) to dark (digit 9). **A**. Tonic–burst with two-stage plasticity (TB_*η*_): weights progressively align with digit samples, showing stable correlation. **B**. Tonic–burst without two-stage plasticity (TB_0_): weights reset at each burst, erasing previously stored information. **C**.Tonic–quiescent model (TQ_0_): quiescence replaces bursts; weights drift and correlations gradually decay. **D**. Tonic-only model (TT_0_): performance fluctuates, with partial correlations but no consolidation across sequences. **E–F**. Generalization percentage during digit classification for sequences I–V, XX, and XXI, measured on **E**. the training set (80 samples) and **F**. an unseen test set (80 samples). TB_*η*_ (dark teal) is generalizing the dataset. TT_0_ (light teal) performs moderately but fails to generalize across sequences. TB_0_ (orange) and TQ_0_ (red) fail to maintain learned paTT_0_erns and show poor generalization. Generalization percentage for TT_0_, TB_0_, and TQ_0_ is normalized to the maximum weight of the TB_*η*_ model for comparison.

Although correlation captures how closely synaptic weights resemble digit templates, it does not directly assess functional performance. To address this, we measure the classification accuracy of the network at the end of each block. This metric reflects the network ability to preserve previously learned information while integrating new inputs. This provides a proxy for consolidation, since high accuracy across sequences indicates retention of prior patterns. We also test performance on a held-out dataset, which quantifies the network capacity to recognize novel inputs based on its acquired representations, *i*.*e*., its ability to generalize.

The accuracy is obtained by testing the network on MNIST digits used during the training protocol under the four models. At the end of each block, the network is presented with 20 samples of each digit from 0 to 3. These samples are either drawn from the training set (used in learning, Figure 2A) or from an unseen test set. This evaluation does not involve synaptic plasticity, allowing us to measure recognition using the learned weights. Predictions are assigned to the output neuron with the highest spike count, and the accuracy is calculated as the fraction of correct predictions. To ensure comparability, models without two-stage plasticity are normalized by the maximum weight value from the TB_*η*_ model (see subsubsection 4.2.9; unnormalized results in Text S4).

On the training set, the TB*η* model shows steadily increasing classification accuracy between sequences and clearly outperforms the other models (Figure 3E). The TB_0_ model shows a sawtooth pattern, with accuracy dropping to near zero after each burst, showing the presence of the burst-induced attractor. The TQ_0_ model learns digits sequentially but fails to maintain them, producing a flat accuracy curve near chance level (25%) before collapsing to zero. The TT_0_ model performs moderately but remains far below TB_*η*_, as each new digit overwrites previous ones. By the final noise sequence, only the TB_*η*_ model maintains robust performance.

On the test set, the TB_*η*_ model also generalizes better: its accuracy increases with time and consistently exceeds the other models (Figure 3F). Although absolute accuracy remains lower than in machine learning systems trained on MNIST, this difference is expected: our goal is not to optimize MNIST performance, but to study how alternating tonic and burst activity supports retention and transfer of information through the two-stage plasticity rule, implemented in biophysical models rather than artificial networks. In this context, accuracy serves as a functional readout of consolidation and generalization mechanisms in a biologically grounded network, rather than as a benchmark of computational efficiency.

### 2.6 Coupling gain and initial state affect consolidation dynamics during burst firing

So far we have presented results using fixed values of the coupling gain 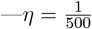 in the small circuit and 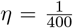 for MNIST—and initial secondary weight, *𝓁*_*ij*_ (0) = 0. 001. In practice, we also observe that consolidation dynamics vary depending on these values. Here, we investigate these effects more systematically in a simplified network. Following the computational protocol of González-Rueda et al. (2018), we assess the information stored in the network across successive states by computing the signal-to-noise ratio (SNR).

We consider a network of 100 presynaptic and 1 postsynaptic neuron (*N* = 100, *M* = 1, Figure 4A), which undergoes 10 alternating states of tonic and burst firing. During each state of tonic firing, 5 randomly selected presynaptic neurons are stimulated by a high-frequency pulse train ranging from 73 to 76 Hz (Figure 4B, dark purple), while the remaining 95 neurons receive stimulation from a low-frequency pulse train ranging from 0.1 to 5 Hz (Figure 4B, lilac). The output neuron is stimulated by a pulse train at an intermediate frequency, fixed at 25 Hz (purple). Synapses undergo the primary and secondary plasticity as introduced in subsection 2.2. During tonic firing, the five neurons firing at a high frequency potentiate their connection with the output neuron, while the rest tend to depress, as dictated by the primary weight calcium-based rule.

**Figure 4:**
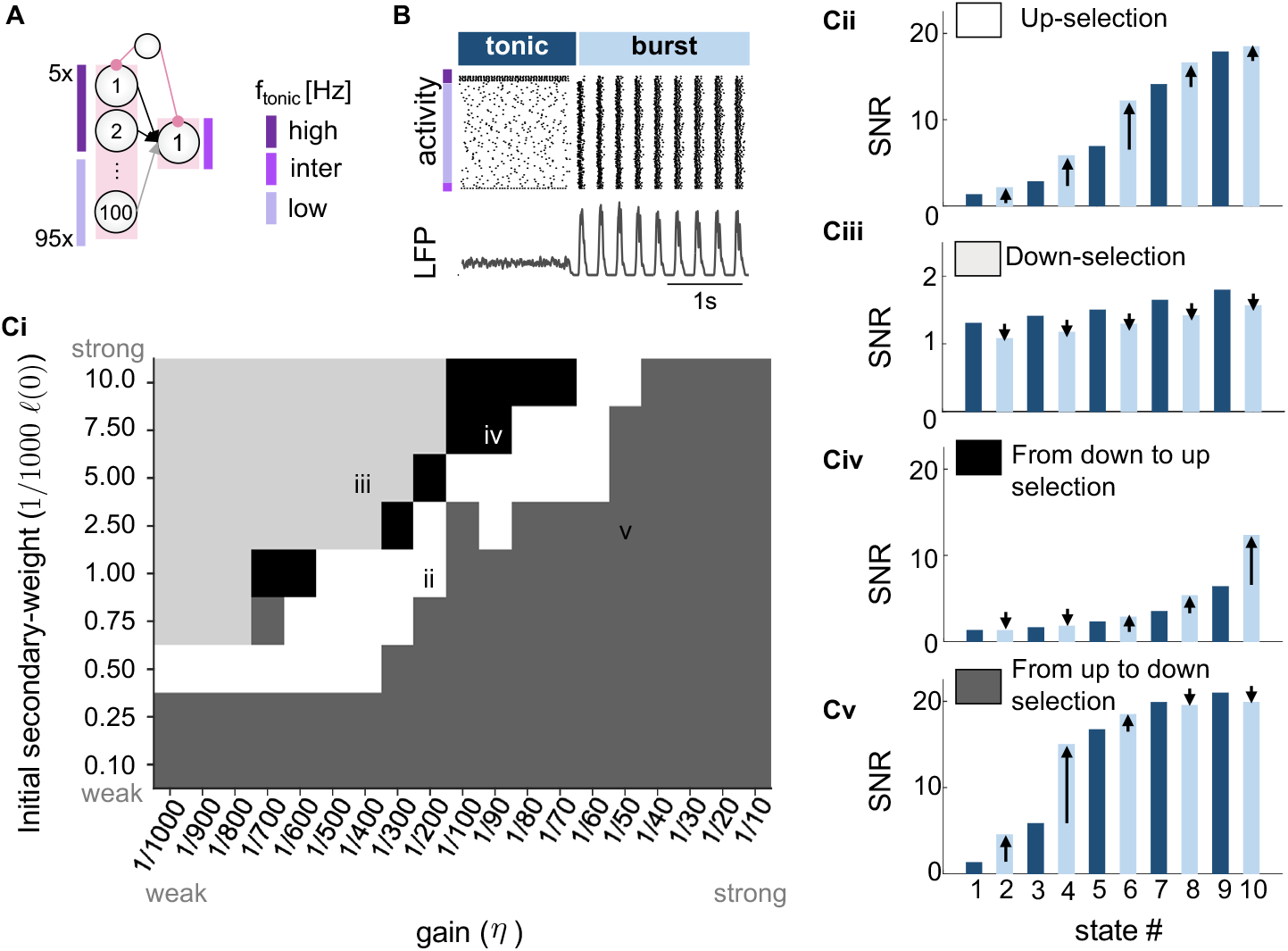
Coupling gain and initial state control synaptic selection during burst firing. **A**. The effect of the coupling gain *η* is studied in a network with one inhibitory neuron projecting to 100 presynaptic neurons connected to a single postsynaptic neuron. The network undergoes 10 cycles of 20 s, alternating between tonic and burst firing (adapted from (González-Rueda et al., 2018)). Presynaptic neurons fire at different tonic frequencies (high, intermediate, or low; shades of purple). **B**. Raster plot of excitatory activity and LFP during the first switch from tonic to burst firing. **C**. Evolution of the signal-to-noise ratio (SNR) across successive tonic and burst states under different parameter settings. Cii. SNR increases during burst firing (up-selection; white, *η* = 1*/*200, *𝓁*_0_ = 0.001). Ciii. SNR decreases during burst firing (down-selection; light gray, *η* = 1*/*400, *𝓁*_0_ = 0.005). Civ. Transition from down-to up-selection during the simulation (black, *η* = 1*/*90, *𝓁*_0_ = 0.0075). Cv. Transition from up-to down-selection (dark gray, *η* = 1*/*50, *𝓁*_0_ = 0.0025).

We define the signal-to-noise ratio (SNR) as the maximum total weight divided by the mean of all total weights (Equation 4) (González-Rueda et al., 2018). Overall, SNR tends to increase over the course of the simulation (Figure 4Cii-v). However, for varying values of *η* and *𝓁*_*ij*_(0), four different microscopic trends emerge across successive tonic and burst firing states (Figure 4Ci). In the first, which we term *up-selection*, the SNR after the burst firing state increases relative to the preceding tonic state (Figure 4Cii). Conversely, during *down-selection*, the SNR decreases during burst firing (Figure 4Ciii). Both mechanisms can coexist: the SNR can initially decrease during burst firing and then increase, in a *transition from down-to up-selection* (Figure 4Civ), as well as the reverse, starting with an increase and then a decrease, in a *transition from up-to down-selection* (Figure 4Cv).

The network can navigate within these four microscopic trends by varying the value of the gain *η*, tuned through the action of neuromodulators, which can influence cascade signaling pathways on a global scale (Brzosko et al., 2019; Pedrosa and Clopath, 2017; Marder et al., 2014; Pawlak and Kerr, 2008; Seol et al., 2007; Lisman et al., 2011) and also the initial secondary weight. For example, dopamine or acetylcholine can affect learning on the whole network (Pedrosa and Clopath, 2017).

Furthermore, these pathways can be locally modulated for specific synapses through synaptic-tagging and capture mechanisms (STC) (Lehr et al., 2022; Frey and Morris, 1997; Ding et al., 2022). According to this theory, synapses tagged after correlated pre- and postsynaptic activity experience enhanced protein synthesis, leading to synapse consolidation.

The effect of the gain, however, varies depending on the initial secondary weight connectivity *𝓁*_*ij*_(0). For low values (*𝓁*_*ij*_(0) ≤ 0.25*e*^*−*3^, the microscopic trend remains the same regardless of the gain (Figure 4Ci, dark grey color). This means that the network relies on burst firing to initially up-select the novel learning and then down-select it, which can be interpreted as a homeostatic technique. However, with intermediate connectivity, there is a range of gain values that promotes up-selection and thus consolidates memory during bursts. For very high connectivity, a weak gain will down-select, a mechanism compatible with synapse pruning that occurs during burst firing (Turrigiano and Nelson, 2004).

In summary, the two-stage synaptic plasticity can lead to both up-selection and down-selection during burst. Up-selection acts as a mechanism for memory consolidation without needing to receive any input. This was also observed in the memorization task (subsection 2.4) through the increased contrast of the synaptic weights during bursting (Figure 2B). Conversely, down-selection prunes previously acquired connections, allowing the network to retain and acquire memory while simultaneously reducing overall connectivity.

### 2.7 Different primary plasticity rules can shape different learning strategies for overlapping patterns

Finally, we examine how the choice of the primary plasticity rule influences memory consolidation in the two-stage synapse model. Although the burst-induced attractor emerges robustly in multiple rules (Jacquerie et al., 2025), the way information is consolidated can differ substantially, particularly when patterns share overlapping features (*i*.*e*., when two images of different categories share common pixels).

We compare two versions of the calcium-based model that governs the primary synaptic plasticity: the model proposed by Graupner and Brunel (2012) (Equation 2) and the one introduced by Graupner et al. (2016) (Equation 3). The 2012 model posits that at low calcium concentrations, the primary weight follows a cubic function with two stable states, while the 2016 model suggests that the primary weight remains constant under low calcium concentrations. In both cases, the secondary plasticity rule remains the same (Equation 1).

To test their impact, we design a simplified recognition task with 16 presynaptic neurons encoding 4 × 4 pixel images and 2 postsynaptic neurons (*N* = 16, *M* = 2, Figure 5A). Each pattern consists of two bars, presented either as non-overlapping (Figure 5B) or overlapping (Figure 5C). Training alternates between tonic and burst states (Figure 5D). During each tonic state, one bar (4 pixels) of the pattern is presented and the subsequent burst state consolidates it before the second bar is introduced. This reflects a scenario where partial features of a class are learned incrementally. We study how the different circuit consolidate the different bars and build the total pattern by studying the weight matrices and the correlation, as it captures how closely synaptic weights resemble to the patterns.

**Figure 5:**
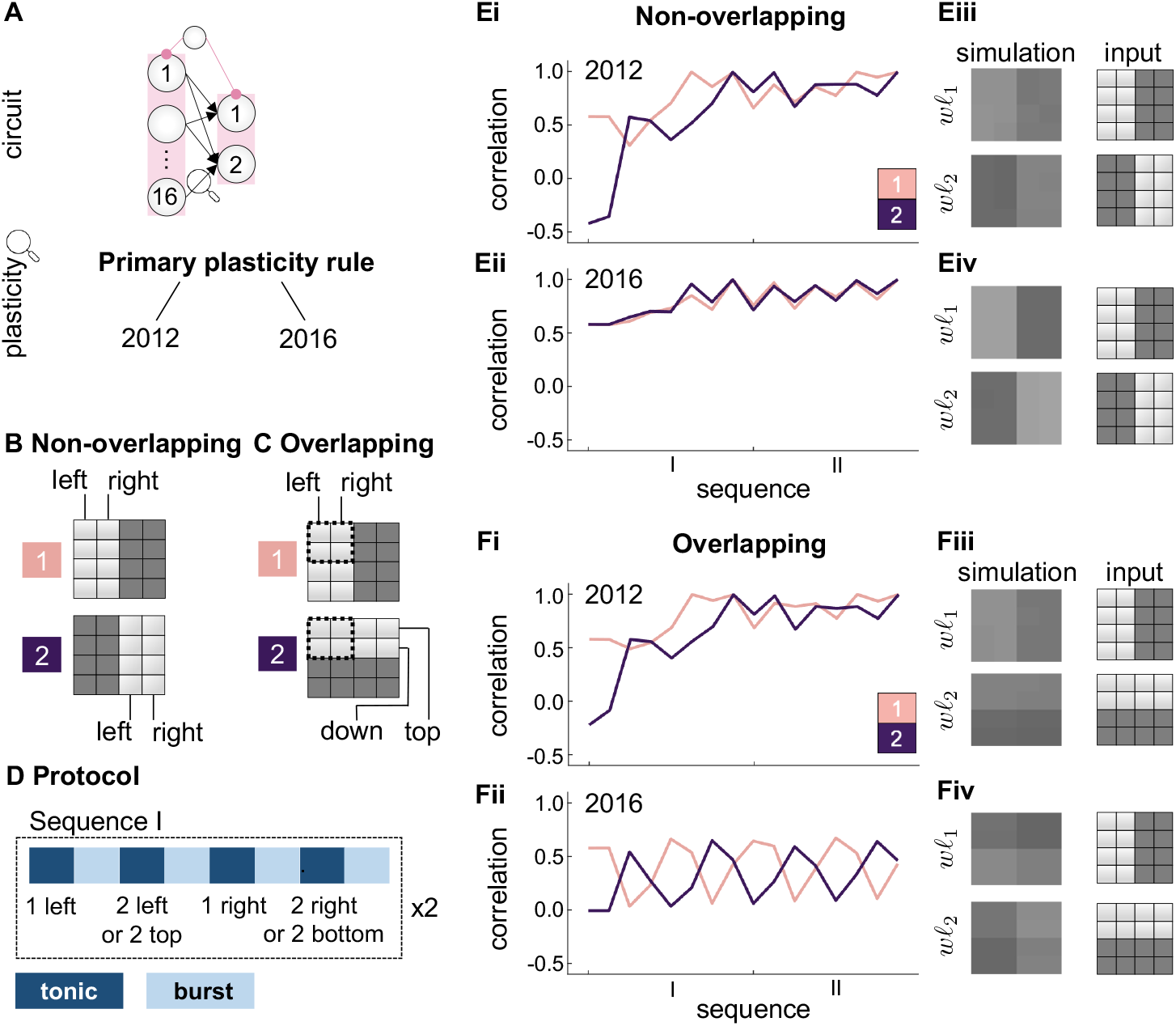
Different primary plasticity rules produce distinct strategies for consolidating overlapping patterns. **A**. Circuit: 16 presynaptic neurons encode 4 × 4 pixel inputs and project to 2 postsynaptic neurons. Excitatory synapses follow either the 2012 calcium-based rule (Graupner and Brunel, 2012) or the 2016 rule (Graupner et al., 2016), combined with secondary plasticity. **B**. The task involves categorizing either non-overlapping patterns (#1, #2) or **C**. overlapping patterns (#1, #2) on a 4×4 grid image. Each pattern is composed of two bars (left, right, or top, down). Shared pixels shown with dashed outline **D**. Training protocol: each sequence alternates tonic (input-driven learning of one bar) and burst states. Each pattern consists of two bars presented sequentially. **E**. Non-overlapping patterns. **(Ei–ii)** Evolution of correlations for the two patterns under the 2012 (Ei) and 2016 (Eii) rules. **(Eiii–iv)** Final weight matrices show successful encoding for both rules, with stronger contrast in the 2016 model. **F**. Overlapping patterns. **(Fi–ii)** Correlations under the 2012 and 2016 rules. The 2012 model consolidates both patterns despite overlap, while the 2016 model suppresses shared pixels. **(Fiii–iv)** Final weight matrices confirm distinct learning strategies.

For non-overlapping patterns, both the 2012 and 2016 models allow successful representations of the patterns in the weight matrices. Correlation traces increase steadily across training sequences (Figure 5Ei–ii), and the final weight matrices match the input patterns (Figure 5Eiii–iv). To quantify selectivity, we compute the contrast between the average synaptic strength of pixels forming the pattern and those outside it. In the 2012 model, contrast is 17% for pattern 1 and 20% for pattern 2, whereas in the 2016 model it rises to 39% for both. Thus, both models support recognition, but the 2016 model yields sharper representations.

For overlapping patterns, the models diverge. With the 2012 rule, both patterns are maintained, as shown by steadily increasing correlations (Figure 5Fi) and final contrasts of 19% and 21%. In contrast, with the 2016 rule, the network fails to reinforce shared pixels: the non-overlapping parts are encoded, but overlapping pixels fade (Figure 5Fii–iv). The resulting contrast drops to 10% for both patterns, and the correlation of one pattern increases only at the expense of the other. The evolution of the weight matrices throughout the protocol in both cases is shown in Figure S3.

These differences arise because the trajectory of the primary weight *w* directly shapes how the secondary weight *𝓁* evolves during bursts. Variations between the 2012 and 2016 calcium-based rules—different parameter values, plasticity constants, or nonlinear terms such as the cubic component in the 2012 formulation—alter the dynamics of *w*, which in turn modifies consolidation through *𝓁*. Although it is difficult to isolate a single factor behind these divergent outcomes, the important point is that our two-stage framework is adaptable: it can be combined with different primary rules, and whichever pathway governs *w* will determine how *𝓁* evolves. In this way, the same mechanism can operate across diverse configurations, producing distinct strategies that favor either integration or separation depending on the underlying primary dynamics.

The 2012 model favors integration, preserving both patterns even when they overlap, while the 2016 model favors separation, encoding only distinct features and discarding shared ones. This resembles how humans sometimes extrapolate from partial cues to reconstruct a full object, yet in other contexts fail to recognize it from fragments alone. In biological circuits, different pathways likely underlie primary plasticity, creating a diversity of consolidation outcomes depending on input structure and context. Our two-stage rule does not interfere with these primary pathways but builds on them, suggesting that biological diversity in plasticity rules may allow circuits to flexibly balance generalization and discrimination according to task demands.

## 3 Discussion

### 3.1 Importance of burst firing in learning

Our model highlights a neuronal firing pattern that has been underexplored in synaptic plasticity research: collective burst firing within a network. Experimental studies typically focus on plasticity under highly controlled conditions, where spikes are precisely timed between presynaptic and postsynaptic neurons (Bi and Poo, 1998; Dudek and Bear, 1992; Inglebert et al., 2020; Sjöström et al., 2001).

In contrast, burst firing emerges from intrinsic neuronal and network mechanisms, and its complexity makes it harder to study experimentally. Computational models often sidestep this by representing rest as quiescence (Fauth and Van Rossum, 2019), yet our results suggest that off-line phase may involve structured burst activity that actively contributes to consolidation.

We show that including burst firing fundamentally changes how memories are stored. Neuronal activity naturally fluctuates between asynchronous (tonic-like) and synchronized (burst-like) regimes, depending on behavioral state and neuromodulation. In our model, bursts provide not just a pause between learning episodes but a distinct phase where synaptic weights are reorganized. The burst-induced attractor pulls synapses to a predictable state, which, when coupled to the secondary weight, stabilizes prior learning. This suggests that alternating between tonic and burst states may be a general strategy for balancing flexibility in encoding with robustness in storage.

### 3.2 Comparison with neo-Hebbian and cascade models

Many plasticity rules have been developed to incorporate additional mechanisms of learning and memory, from eligibility traces to cascade models. Our two-stage rule shares some of their goals but differs in key ways.

Eligibility-trace models compute a decaying variable from pre- and postsynaptic activity, which only influences plasticity when gated by a third factor such as reward or neuromodulation (Gerstner et al., 2018). In our framework, no extra variable is needed: bursting itself provides a built-in trace, as the drift of the primary weight toward the attractor is transferred into the secondary weight 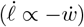. Consolidation is therefore tied to network state rather than to an external signal, and the effective weight is defined by the multiplicative interaction of the two variables.

Cascade models instead assign each synapse a hierarchy of hidden states with different stabilities, where plasticity events trigger transitions and memory lifetimes emerge from slow diffusion across states (Fusi et al., 2005). In contrast, our rule uses only two variables and one coupling parameter, exploiting the burst-induced attractor as a built-in trace for consolidation, without requiring multiple hidden states or stochastic transitions.

These frameworks should not be viewed as mutually exclusive. Future models may combine elements of cascade-like stabilization, eligibility traces, and state-dependent mechanisms such as ours, depending on the computational demands of the task (Zenke et al., 2015).

### 3.3 Implications for spiking neural networks and neuromorphic design

Beyond biological circuits, our findings also carry implications for artificial and neuromorphic networks. Most current spiking models rely exclusively on tonic spiking to represent and learn information. Here, alternating tonic and burst phases offer a new route for synaptic reorganization and consolidation (Hinton et al., 1995; Legenstein and Maass, 2007). The burst state acts as an “internal consolidation phase,” during which information is stabilized without replay of prior inputs. This provides a biologically inspired mechanism that could be integrated into neuromorphic systems to improve learning.

More generally, the fluctuating activity of biological circuits points to a useful design principle: networks should be trained not in a single spiking regime but with structured alternations between input-driven tonic activity and collective bursting modes. Our two-stage rule is particularly well suited to such applications. The effective weight is the product of two variables, with the secondary updated by a local derivative-based term. The coupling gain can be tuned as a single parameter, making the mechanism computationally lightweight and straightforward to implement in artificial networks or neuromorphic frameworks.

### 3.4 Two-stage plasticity: simplicity, potential, and limitations

Finally, we place our approach in the broader landscape of plasticity rules, which span a wide spectrum—from short-term plasticity to structural plasticity, from detailed biophysical models of protein kinetics (Smolen et al., 2006) to abstract spike-based correlation rules (Deger et al., 2012). The former provide mechanistic insight but are computationally expensive, while the latter are analytically tractable but often restricted to tonic activity, making them poorly suited for bursting dynamics.

An alternative framework for consolidation is synaptic tagging and capture (STC) (Seibt and Frank, 2019; Smolen et al., 2020), where potentiated synapses are locally tagged and stabilized through plasticity-related proteins. In models, STC is often implemented by summing primary and secondary weights (Luboeinski and Tetzlaff, 2021; Clopath et al., 2008). While this additive formulation works under tonic learning, it fails during bursting: the burst-induced attractor drags all primary weights in the same direction, pulling the secondary weights with them.

The two-stage rule we propose offers a simpler and more robust alternative. It requires only one equation with one free parameter (the coupling gain *η*), yet consolidates memory by exploiting—rather than being undermined by—the burst-induced attractor. In this framework, the attractor becomes a built-in trace that the secondary weight can use to stabilize previously acquired learning.

This simplicity opens multiple avenues for extension. First, we assume secondary plasticity is only active during bursts (*η* = 0 in tonic, *η >* 0 in bursts). In reality, signaling pathways may also occur during tonic activity; allowing a small negative *η* could let the secondary weight accumulate alongside primary potentiation, priming synapses for later consolidation. Second, we modeled *η* as uniform, but in biological systems it could vary between synapses depending on activity tags or neuromodulatory signals. Introducing heterogeneous *η* would enable selective consolidation of salient memories. Third, the coupling is modeled as a simple linear dependence on the derivative of *w*. More complex couplings—such as making *η* depend on weight history or total strength—could prevent excessive growth and improve stability. Exploring such refinements may bring the model closer to biological reality and broaden its applicability to more complex tasks.

## 4 METHOD

### 4.1 Data and code availability

All original data in this work were generated programmatically using the Julia programming language (Bezanson et al., 2017). Analyses were performed in Matlab. The code files are freely available at https://github.com/KJacquerie/Two-stage-Plasticity. Any additional information required to reanalyze the data reported in this paper is available from the lead contact upon request.

### 4.2 Method details

#### 4.2.1 Neuron and network model

All neurons are modeled using a single-compartment conductance-based using Hodgkin and Huxley formalism (Hodgkin and Huxley, 1952). The membrane voltage *V* of a neuron evolves according to the equation described by (Drion et al., 2018; Jacquerie and Drion, 2021):

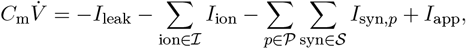

where *C*_m_ represents the membrane capacitance, *I*_leak_ is the leak current, *I*_ion_ are the intrinsic ionic currents with ℐ the set of all ionic channels, *I*_syn,*p*_ are the synaptic currents with 𝒮 the set of all synaptic neurotransmitter types and 𝒫 the set of all presynaptic neurons, and *I*_app_ denotes the applied current. Details about the ionic and synaptic currents and their associated dynamics and parameters are provided in (Jacquerie et al., 2025).

The neuronal network comprises an inhibitory neuron (𝒩_inh_) projecting onto all excitatory neurons through GABA_A_ and GABA_B_ connections (refer to Figure 1B). Excitatory neurons are interconnected via a feedforward AMPA synapse, where the presynaptic neurons (𝒩_pre_) influence the postsynaptic neurons (𝒩_post_). The number of excitatory neurons varies across different computational experiments. The excitatory synaptic current perceived by the postsynaptic neuron *i* from presynaptic neuron *j* is characterized by:

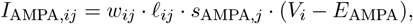

where *w*_*ij*_ represents the primary weight, *𝓁*_*ij*_ is the secondary weight, and their product *w*_*ij*_ *· 𝓁*_*ij*_ defines the total synaptic weight. The variable *s*_AMPA,*j*_ denotes the gating variable of the AMPA postsynaptic receptor (AMPAr), dynamically modulated by the presynaptic membrane voltage (*V*_*j*_) and *E*_AMPA_ is the reversal potential of AMPAr. This model extends our previous representation of 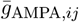 as the maximal conductance of the AMPA receptors to accommodate primary and secondary synaptic plasticity, as outlined in (Jacquerie and Drion, 2021).

#### 4.2.2 Switch from tonic to burst firing

We follow the same method as in (Jacquerie et al., 2025). The inhibitory neuron dynamically influences network activity. An external current applied to the inhibitory neuron (*I*_app,inh_) models the effects of a neuromodulatory signal. A depolarizing current induces tonic firing activity in this inhibitory neuron, causing all excitatory cells to remain at rest. A sufficiently large external depolarizing pulse can evoke action potentials in excitatory neurons. Conversely, a hyperpolarizing current applied to the inhibitory neuron switches the entire network into a synchronized collective bursting activity throughout the network (Zagha and McCormick, 2014; Jacquerie and Drion, 2021). In each computational experiment, the depolarizing current *I*_app,inh_ is equal to 3 nA*/*cm^2^ for tonic state and −1.2 nA*/*cm^2^ for burst state.

#### 4.2.3 External applied pulse train current

We follow the same method as in (Jacquerie et al., 2025). In either tonic firing, quiescent states, or noisy states, each excitatory neuron *i* is triggered by an applied current (*I*_app,*i*_) to make the neuron fire at a nominal frequency *f*_0_. To generate this input-driven tonic activity, a neuron receives a pulse train current, where each pulse lasts 3 ms and has an amplitude that is independently sampled from a uniform distribution on an interval between 50 nA*/*cm^2^ and 60 nA*/*cm^2^. The interpulse intervals (*i*.*e*., the time between two successive pulses) are independently sampled from a Normal distribution with a mean equal to 1*/f*_0_ and a standard deviation equal to 0.1*/f*_0_.

In burst firing states, each excitatory neuron *i* undergoes a noisy baseline such as the applied current (*I*_app,*i*_) randomly fluctuating between the interval 0 and 1 nA*/*cm^2^.

#### 4.2.4 Homogeneous and heterogeneous network

We follow the same method as in (Jacquerie et al., 2025). We define a *homogeneous* network (or circuit), where each neuron has the same intrinsic ion channel maximal conductances 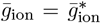 (numerical values are provided in subsubsection 4.2.4). We define a *heterogeneous* network (or circuit), where the maximal conductance 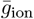 for each neuron is randomly chosen within a ±10 % interval around its nominal value 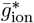, such as 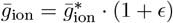 with *ϵ* ∼ Unif(−0.1, 0.1).

#### 4.2.5 Local field potential

We follow the same method as in (Jacquerie et al., 2025). The local field potential (LFP) measures the average behavior of interacting neurons. It reflects the collective excitatory synaptic activity received by the postsynaptic neuron population. The overall synaptic activity is measured by the mean of the individual synaptic currents:

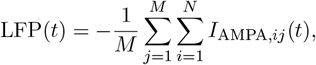

where *M* is the number of postsynaptic neurons and *N* is the number of presynaptic neurons (Jacquerie and Drion, 2021; Drion et al., 2018).

#### 4.2.6 Synaptic plasticity

##### Primary synaptic plasticity

The change in the primary weight *w*_*ij*_ between a presynaptic neuron *j* and a postsynaptic neuron *i* is governed by the calcium-based model proposed by (Graupner and Brunel, 2012):

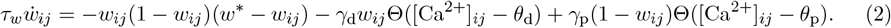

Here, *τ*_*w*_ represents the time constant, *w*^*\**^ defines the stable state (equal to 0.5), *γ*_p_ is the potentiation rate, *γ*_d_ is the depression rate, *θ*_p_ is the potentiation threshold, and *θ*_d_ is the depression threshold. The function Θ(·) is the Heaviside function, which returns 1 if [Ca^2+^]_*ij*_ *> θ*_p_ and 0 otherwise.

The change in primary weight depends on the calcium concentration [Ca^2+^]_*ij*_, which is the sum of the calcium caused by the activity of the presynaptic neuron *j* and the activity of the postsynaptic neuron *i*. A pre- or postsynaptic spike translates into a calcium exponential decay. For details about the calcium dynamics, see Text S1. The calcium-based rule defined by (Graupner and Brunel, 2012) implements a soft-bound, where the perceived potentiation rate *γ*_p_(1 − *w*_*ij*_) is smaller for high *w*_*ij*_ than for low *w*_*ij*_. The same reasoning applies to the depression rate *γ*_d_. Detailed parameter values are provided in Text S2.

In Figure 5, the calcium-based model is modified according to the version of (Graupner et al., 2016). The model is similar except that the stable state is removed and the fitted plasticity parameters are adapted:

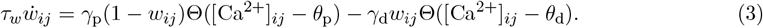

##### Secondary synaptic plasticity

The secondary weight *𝓁*_*ij*_ between the presynaptic neuron *j* and the postsynaptic neuron *i* evolves according to Equation 1. The coupling gain *η* is set to 1/500 in Figure 1, 1/400 in Figure 2, varies from 1/1000 to 1/10 in Figure 4, and 1/200 in Figure 5.

The threshold *θ*_thres_ was not assigned a single fixed value but adjusted to ensure clear separation between tonic and burst regimes in each network instantiation. In practice, tonic firing emerges for *I*_app,inh_ ≳ 0, while collective bursting is robust for *I*_app,inh_ ≲ −1. Exact values are not critical, reflecting the high degree of degeneracy typical of biophysical networks.

#### 4.2.7 Computational experiment related to Figure 1

Figure 1C illustrates the activity in the feedforward network, comprising 50 presynaptic neurons connected to 50 postsynaptic neurons. The network is heterogeneous. The network is in tonic firing mode during 1.5 s (see subsubsection 4.2.2). Each excitatory neuron receives a train of current pulses with nominal frequencies *f*_0_ randomly sampled from a uniform distribution between 0.1 and 50 Hz (see subsubsection 4.2.3). Then, the network switches to synchronized collective bursting during 2.5 s.

Figure 1E–F show results from a small heterogeneous circuit comprising one inhibitory neuron providing GABAergic input to four excitatory neurons, where two presynaptic excitatory neurons project onto two postsynaptic excitatory neurons.

The computational experiments include four configurations: alternating tonic and burst states (Figure 1D-E, tonic states interleaved with quiescent periods (Figure 1F), or continuous tonic states (Figure 1G). In Figure 1D, primary and secondary plasticity are active, while in Figure 1E both the secondary plasticity is blocked 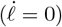.

During tonic states, excitatory neurons are stimulated with pulse trains at fixed nominal frequencies. In the first tonic state, the two presynaptic neurons fire at 60 Hz and 1 Hz, and the postsynaptic neurons at 35 Hz and 5 Hz. In the second tonic state, presynaptic neurons fire at 65 Hz and 1 Hz, while postsynaptic neurons fire at 30 Hz and 5 Hz. These values were selected to illustrate potentiation and depression during tonic activity. Details of the current injection and interpulse intervals are provided in subsubsection 4.2.3.

During burst states, the inhibitory neuron is hyperpolarized, producing collective burst firing as described in subsubsection 4.2.2. In the quiescent condition, all neurons are driven by low-frequency pulse trains randomly selected from 0.1 Hz, 0.5 Hz, or 1 Hz.

In the additional tonic condition (Figure 1G), firing frequencies are randomly drawn from the set {0.5, 1, 5, 10, 40, 50} Hz.

#### 4.2.8 Computational experiment related to Figure 2

Figure 2A illustrates the simulation protocol for a learning task interleaved with periods of burst firing. The whole task lasts 6030 s (*i*.*e*., 100 min and 30 s). The protocol is divided into 21 sequences (Roman numerals). In each sequence (I-XX), the network sequentially learns the digits 0 to 9. Each sequence lasts a total of 300 s and is composed of 10 blocks of tonic and burst firing. Each block lasts 30 s with a tonic or a burst firing state of 15 s each. The final sequence (XXI) lasts 30 s and consists of only one block. A cropped version of the MNIST dataset (LeCun et al., 1998) with 22 ×22 pixels is used. The initial dataset of 28 × 28 pixels has 3 pixels cropped on each side. The training set is composed of 20 samples of each digits pulled in the training set composed of 80 samples.

The network consists of 484 (22 × 22) presynaptic neurons representing individual image pixels and 10 postsynaptic neurons representing the digit classes (0 to 9). One inhibitory neuron projects onto all these neurons. The network is homogeneous to eliminate confounding factors while studying variations in the synaptic plasticity rule (parameters are available in (Jacquerie et al., 2025)).

A learning state in the network corresponds to tonic activity (dark blue). Excitatory neurons are initially at resting membrane potential. Presynaptic neurons are activated by a train of current pulses with a nominal frequency reflecting the pixel color. The MNIST images are binarized such that a pixel with an intensity greater than 0.5 is considered a white pixel otherwise it is black. A white pixel is encoded by a neuron spiking at 45 Hz and a black pixel at 0.01 Hz. The postsynaptic neuron corresponding to the presented digit class spikes at 45 Hz, while the others spike at 0.0001 Hz. Each sequence is composed of 10 blocks. The learning states are sequentially organized from digit 0 to digit 9, with one sample digit randomly chosen from the MNIST dataset for each digit class.

An off state corresponds to either collective bursting activity (light blue) or quiescent activity (lilac) in the network. For the collective bursting activity, all excitatory neurons do not receive any input-driven pulse but only undergo baseline noise (subsubsection 4.2.3). For the quiescent activity, all neurons receive a pulse train with nominal frequencies independently sampled from a Normal distribution with a mean equal to 1 Hz and a standard deviation equal to 0.1 Hz.

A noisy state (gray) occurs during sequence XXI for 15 s. In this state, all excitatory neurons receive a pulse train with nominal frequencies independently sampled from a Normal distribution with a mean equal to 15 Hz and a standard deviation equal to 5 Hz.

Figure 2B-E illustrates the weight matrix of each digit. It corresponds to the total synaptic weights of the 482 presynaptic neurons onto the postsynaptic neuron associated with the considered digit, reshaped in a 22 × 22 grid matrix. The color of each pixel is proportional to the synaptic weight matrix (*w𝓁*)_*i*_ acquired at the end of each sequence (shown in Figure S1). For visualization purposes, synaptic weights are normalized so that the normalized synaptic weight values throughout the whole simulation span the interval (0, 1), that is,

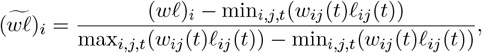

where the maximum and minimum values are obtained by comparing the values across all pairs of presynaptic and postsynaptic neurons at the end of each state throughout the whole simulation. The weight matrices are shown for the states 1, 2 (in sequence I), 201, 202, 239, 240 (in sequence XI), 399, 400 (in sequence XX), 401, and 402 (in sequence XXI) for the digit 0 and 9. Other digits can be seen in Figure S1.

Figure 2B alternates between tonic and burst firing with the two-stage plasticity rule (*η* = 1*/*400). Figure 2C blocks the secondary synaptic plasticity with a coupling gain of 0 in burst firing too. Figure 2D replace the bursting state by a quiescent state. Figure 2E, except that the bursting state (light blue) is replaced by an additional learning state for the same digit, using a different randomly chosen sample of the same digit. The weight matrices are obtained in the same manner, and the normalization is achieved by selecting the maximum and minimum values in this simulation. They differ from those in Figure 2B (see Text S2 for the numeric values).

#### 4.2.9 Computational experiment related to Figure 3

Figure 3A-D illustrate the evolution of the two-dimensional correlation coefficient between the mean image of the dataset and the resultant normalized weight matrix at the end of each state, as obtained from the task depicted in Figure 2B-E. The mean image of the dataset is computed by averaging the pixel values across the 20 images per digit used to train the network (with one image per sequence) (see last column Figure 2B,D-E). ForTT_0_ model, the mean image of the dataset employed in Figure 2E consists of 40 images instead of 20.

Figure 3E-F show the evolution of consolidation percentage at each state of the learning task. Eighty samples are presented to the network, consisting of 20 samples from digit classes 0 to 3. Samples are either selected from the training set (Figure 3E) or the testing set (Figure 3F) and processed according to the methods described in subsubsection 4.2.8. Each sample is presented for 2 s. The external current applied to the inhibitory neuron is set to 0 nA*/*cm^2^ to modulate the network overall excitability, mimicking a retrieval state. Presynaptic neurons are activated by a train of current pulses, with the frequency reflecting pixel intensity: white pixels are encoded by a neuron spiking at 55 Hz, and black pixels at 0.0001 Hz. Postsynaptic neurons are depolarized by 6.5 nA*/*cm^2^ and spike at 0.0001 Hz.

Plasticity is turned off, and synaptic weights are frozen at the values obtained at the end of each state (unnormalized weights). Since secondary plasticity is only active in the TB_*η*_ model, we also consider normalized weights at a given state *T*^*\**^ for theTT_0_, TB_0_, and TQ_0_ models. The normalized weights are calculated as follows:

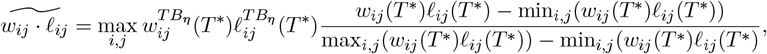

where 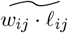 represents the normalized total weight, 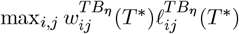 is the scaling factor corresponding to the maximum weights at state *T*^*\**^ in the TB_*η*_ model, and the minimum and maximum values are taken across all presynaptic and postsynaptic neuron pairs at the end of state *T*^*\**^ for the respective model (TT_0_, TB_0_, or TQ_0_).

The number of spikes generated by each output neuron is counted after a transient period of 0.5 s. The network prediction is determined by the output neuron with the highest spike count. Consolidation percentage is defined as the percentage of correct predictions out of the 80 samples tested.

#### 4.2.10 Computational experiment related to Figure 4

Figure 4A illustrates a heterogeneous network comprising one inhibitory neuron connected to 101 excitatory neurons, where 100 presynaptic neurons synapse onto 1 postsynaptic neuron. The stimulation protocol consists of 10 states interleaving tonic firing and burst firing, each lasting 15 s.

For each tonic state, the first five neurons (dark purple) receive an applied current that consists of a train of pulses with a frequency independently sampled for each neuron from a uniform distribution between 73 and 76 Hz (the frequencies are resampled at the beginning of each tonic state). The other 95 presynaptic neurons (lilac) receive a pulse train with a frequency independently sampled for each neuron from a uniform distribution between 0.1 and 5 Hz. The postsynaptic neurons receive a pulse train at 25 Hz (purple). These values are selected to demonstrate strong and weak correlations, although other ranges of values also yield results. The pulse train is constructed as previously described. The raster plot and the LFP are computed for 4 s during the first state transition (Figure 4B). In Figure 4C, the signal-to-noise ratio is computed following the method proposed by (González-Rueda et al., 2018):

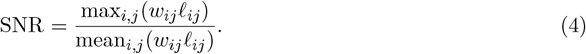

Figure 4Ci highlights the trend followed by the SNR for different values of the coupling gain (*η*) and the initial value of the secondary weight (*𝓁*_0_). Figure 4Cii to v are obtained for the following values: (ii) *η* = 1*/*200, *𝓁*_0_ = 0.001; (iii) *η* = 1*/*400, *𝓁*_0_ = 0.005; (iv) *η* = 1*/*90, *𝓁*_0_ = 0.0075; (v) *η* = 1*/*50, *𝓁*_0_ = 0.0025.

#### 4.2.11 Computational experiment related to Figure 5

Figure 5A illustrates a homogeneous network comprising one inhibitory neuron connected to 18 excitatory neurons, where 16 presynaptic neurons synapse onto 2 postsynaptic neurons. Synapses undergo synaptic plasticity where the primary weight is either driven by the primary plasticity fitted in the calcium-based rule of (Graupner and Brunel, 2012) (2012 model) or (Graupner et al., 2016) (2016 model). The secondary plasticity is active across both scenarios. Detailed parameters can be found in subsection 4.2.

The network is trained to learn two patterns, each corresponding to 8 active pixels out of the 16pixel grid. The simulation protocol consists of two sequences of 4 tonic firing states interleaved with four burst firing states. During each tonic state, only 4 out of the 8 pixels are learned. It corresponds to creating samples for the same class. To achieve this, the corresponding input neurons are activated by a pulse train at 40 Hz, while the output neuron associated with the pattern class is triggered by a pulse train at 55 Hz. Inactive input pixels receive a pulse train at 0.01 Hz, and the second output neuron receives a pulse train at 0.0001 Hz. We alternate between learning one part of a pattern and, in the subsequent tonic firing state, one part of the other pattern. Correlation is computed as detailed in the methods section for Figure 2.

To compute the percentage difference, *µ*_in_ is defined as the mean of the pixels within the pattern, and *µ*_out_ as the mean of the pixels outside the pattern. In the non-overlapping case, for the first pattern, *µ*_in_ is the mean of the first two columns, and *µ*_out_ is the mean of the third and fourth columns. For the second pattern, these are reversed. In the overlapping case, *µ*_in_ and *µ*_out_ are calculated similarly, except for the second pattern, where *µ*_in_ is the mean of the first two rows, and *µ*_out_ is the mean of the third and fourth rows. The percentage difference is defined as (*µ*_in_ − *µ*_out_)*/*((*µ*_in_ + *µ*_out_)*/*2).

## Acknowledgments

Kathleen Jacquerie was a Research Fellow of the Fonds de la Recherche Scientifique—FNRS. This work was supported by the Belgian Government through the Federal Public Service Policy and Support. The authors thank Juliette Ponnet, Nora Benghalem, Justine Magis, and Emmy Kellens for their contributions during their Master’s theses, as well as Caroline Minne for her involvement in the early discussions that shaped the conceptualization of this project. We are also grateful to Professor Eve Marder for her valuable insights and feedback.

## Supplementary Material

### Text S1 Calcium dynamics

We consider the linear calcium dynamics suggested by (Graupner and Brunel, 2012; Graupner et al., 2016). The presynaptic and postsynaptic spike contributions add linearly. Indeed, calcium enters from NMDA receptors and voltage-dependent calcium channels. Instead of describing the whole calcium, the phenomenological effect on the calcium variation is considered. At each pre or postsynaptic spike (respectively named as *t*_*j,k*_ and *t*_*i,k*_), the calcium immediately rises and then exponentially decays characterized by a calcium decay time constant equal to *τ*_Ca_:

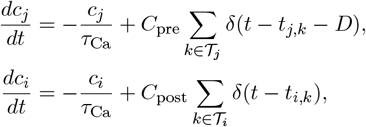

where *C*_pre_ and *C*_post_ are the presynaptically and postsynaptically evoked calcium amplitudes. The parameter *D* is a time-delay between the presynaptic spike and the corresponding postsynaptic calcium transient occurrence accounts for the slow rise time of the NMDAr-mediated calcium influx (Graupner and Brunel, 2012; Graupner et al., 2016).

The total calcium amplitude [Ca^2+^]_*ij*_(*t*) driving the synaptic change is given by:

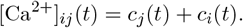

The time-evolution for several pre- and postsynaptic spiking activity is written such as (Graupner et al., 2016):

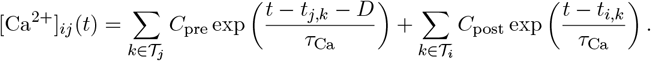

The resting calcium concentration is set to zero. The calcium concentrations are dimensionless. Both simplification is acknowledged because the synaptic rules are adapted in accordance. If a resting calcium concentration is wanted, the thresholds of potentiation and depression will be adapted. This notation follows the original paper notation.

### Text S2 Computational experiments

The parameters associated with the primary synaptic plasticity are given in (Graupner and Brunel, 2012). *τ*_Ca_ = 22.6936 ms, *C*_pre_ = 0.56, *C*_post_ = 1.24, *D* = 4.60 ms, *τ*_*w*_ = 346.3615 ×10^3^ ms, *γ*_p_ = 725.085 × 1.1 (Tonic), *γ*_p_ = 725.085 ×0.95 (Burst), *γ*_d_ = 331.909, *θ*_p_ = 1.3, *θ*_d_ = 1, *w*^*\**^ = 0.5. The potentiation rate *γ*_*p*_ is slightly scaled up during tonic firing to induce stronger potentiation compared to the initial model, and it is reduced by 5 % during burst firing to place the reset at a lower value compared to the initial model.

The parameters used in each simulation are *N* = 50, *M* = 50, *T*_state_ = 20 s, *N*_state_ = 8 for Figure 1, *N* = 484, *M* = 10, *T*_state_ = 15 s, *N*_state_ = 62 for Figure 2 and Figure 3, *N* = 50, *M* = 50, *T*_state_ = 20 s, *N*_state_ = 4 for Figure 4, *N* = 100, *M* = 1, *T*_state_ = 20 s, *N*_state_ = 10 for Figure 4, and *N* = 16, *M* = 2, *T*_state_ = 15 s, *N*_state_ = 16 for Figure 5. Some parameters remain constant in the different simulations: *I*_app,inh_(Tonic) = 3 nA*/*cm^2^, *I*_app,inh_ (Burst) = −1.2 nA*/*cm^2^, *w*_0_=0.5, *𝓁*_0_=0.001, *η* = 1/500 in Figure 1, 1/400 in Figure 2, and 1/200 in Figure 5. In Figure 4, *𝓁*_0_ varies from 0.0001 to 0.01 and *η* varies from 1*/*1000 to 1*/*10. In Figure 5, *𝓁*_0_ equals 0.02. To constrain the secondary weight, a maximum upper bound is present (*𝓁*_max_) - this bound is never reached in the different computational experiments. The minimum and maximum values of *w𝓁* are, respectively, 0.0265*e*^*−*3^, 0.00342 for Figure 2B, 0.168*e*^*−*3^, 0.733*e*^*−*3^ for Figure 2C, and 0.176*e*^*−*3^, 0.706*e*^*−*3^ for Figure 4A-B.

### Text S3 Supplementary material related to Figure 2

Figure S1 shows the evolution of the weight matrices associated with Figure 2.

**Figure S1:**
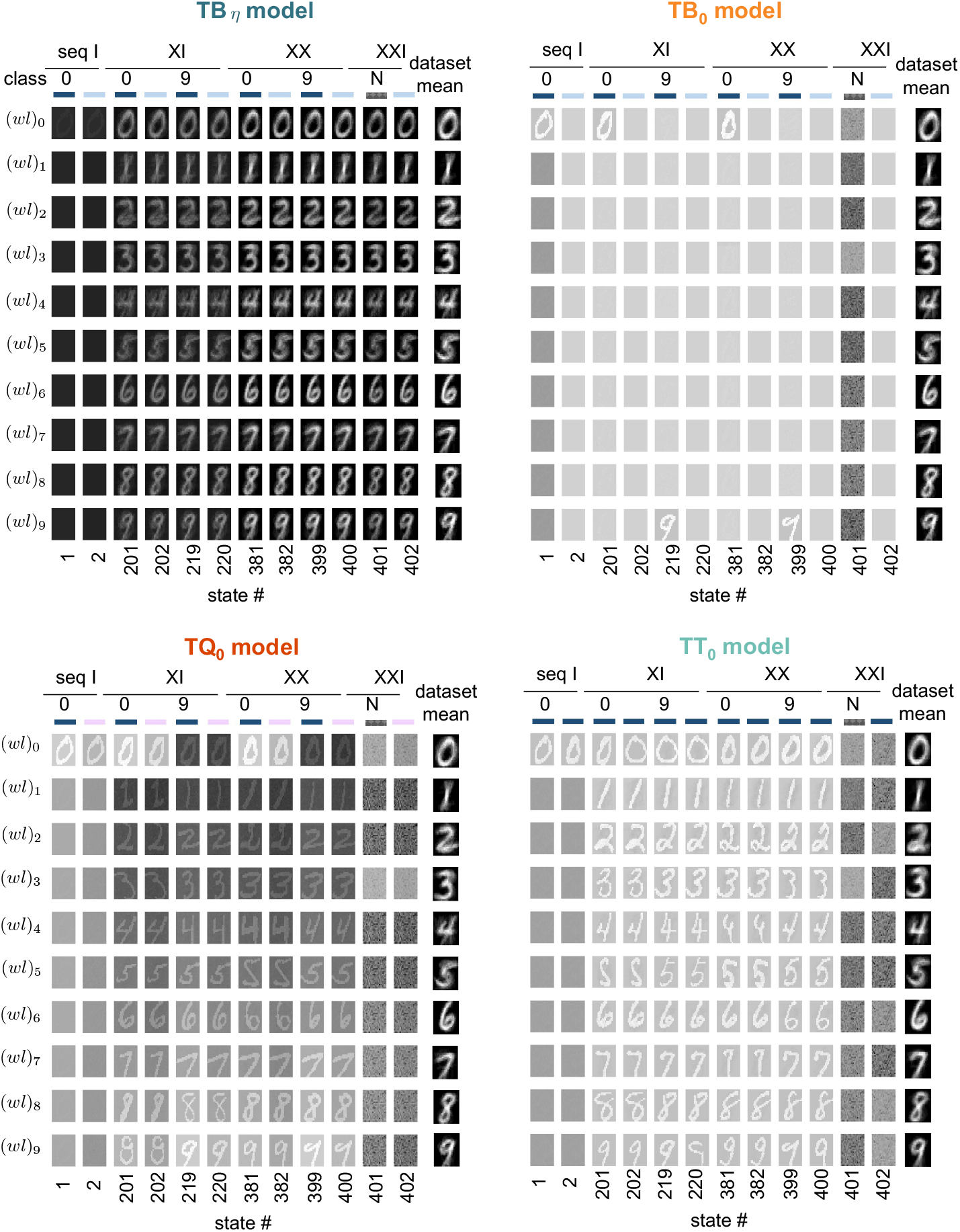
Evolution of the weight matrices for the 10 digits at different states, associated with Figure 2 for the different models (TB_*η*_, TB_0_, TQ_0_,TT_0_), depicts the protocol that interleaves different firing activities depending on the model color coded by tonic (dark blue), burst firing (light blue), noise (for N, in gray) and quiescent (lilac). (seq means sequence).

### Text S4 Supplementary material related to Figure 3

We compare the effect of normalization on training and testing consolidation percentage. Figure S2A-B show results for unnormalized weights, while Figure S2C-D display normalized weights. Testing involves unseen samples of digits 0 to 3. Models without secondary plasticity, where only primary weights fluctuate, perform poorly, while our TB_*η*_ model excels in the training set and can generalize its learning on the testing set. Normalization improves the performance of other models but still lags behind our approach.

**Figure S2:**
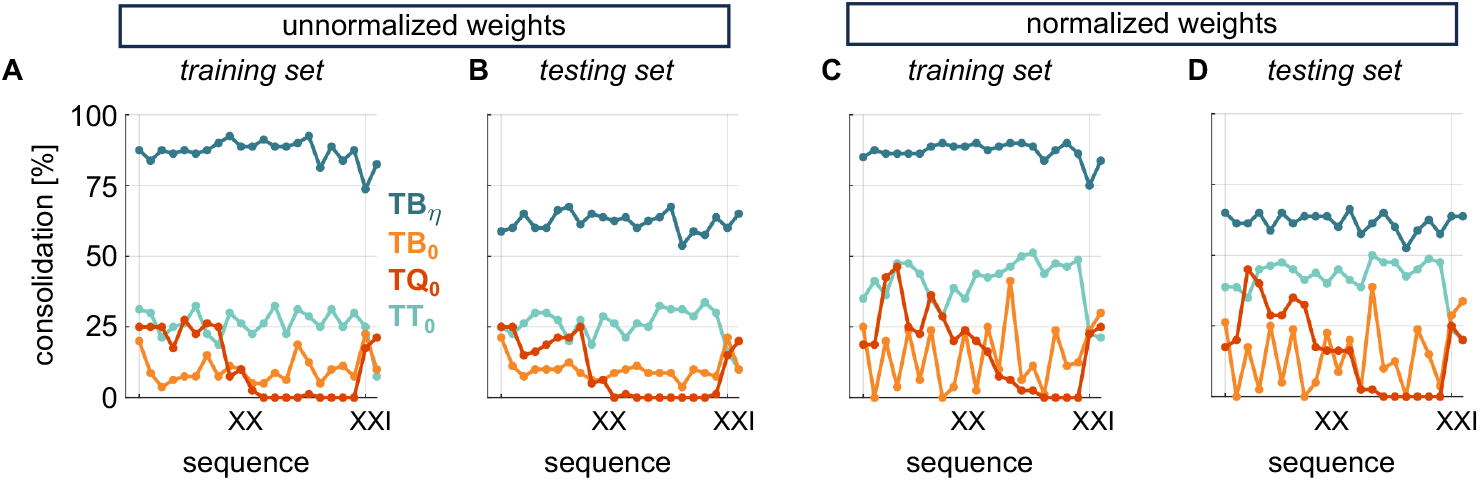
Comparison of weight normalization on consolidation percentage for training and testing datasets. Consolidation percentage was obtained for the final two sequences (XX-XXI) without weight normalization**(A, B.)**, compared to with normalized weights **(C, D)**. Consolidation percentage is compared between the training dataset **(A, C)** and the testing dataset **(B, D)**.

### Text S5 Supplementary material related to Figure 5

Figure S3 depicts the evolution of the weight matrices associated with Figure 5 across successive tonic and burst firing states. In Figure S3A, the weight matrices are compared for non-overlapping conditions between the 2012 model and the 2016 model. Similarly, Figure S3B compares the weight matrices under overlapping conditions between the 2012 model and the 2016 model.

**Figure S3:**
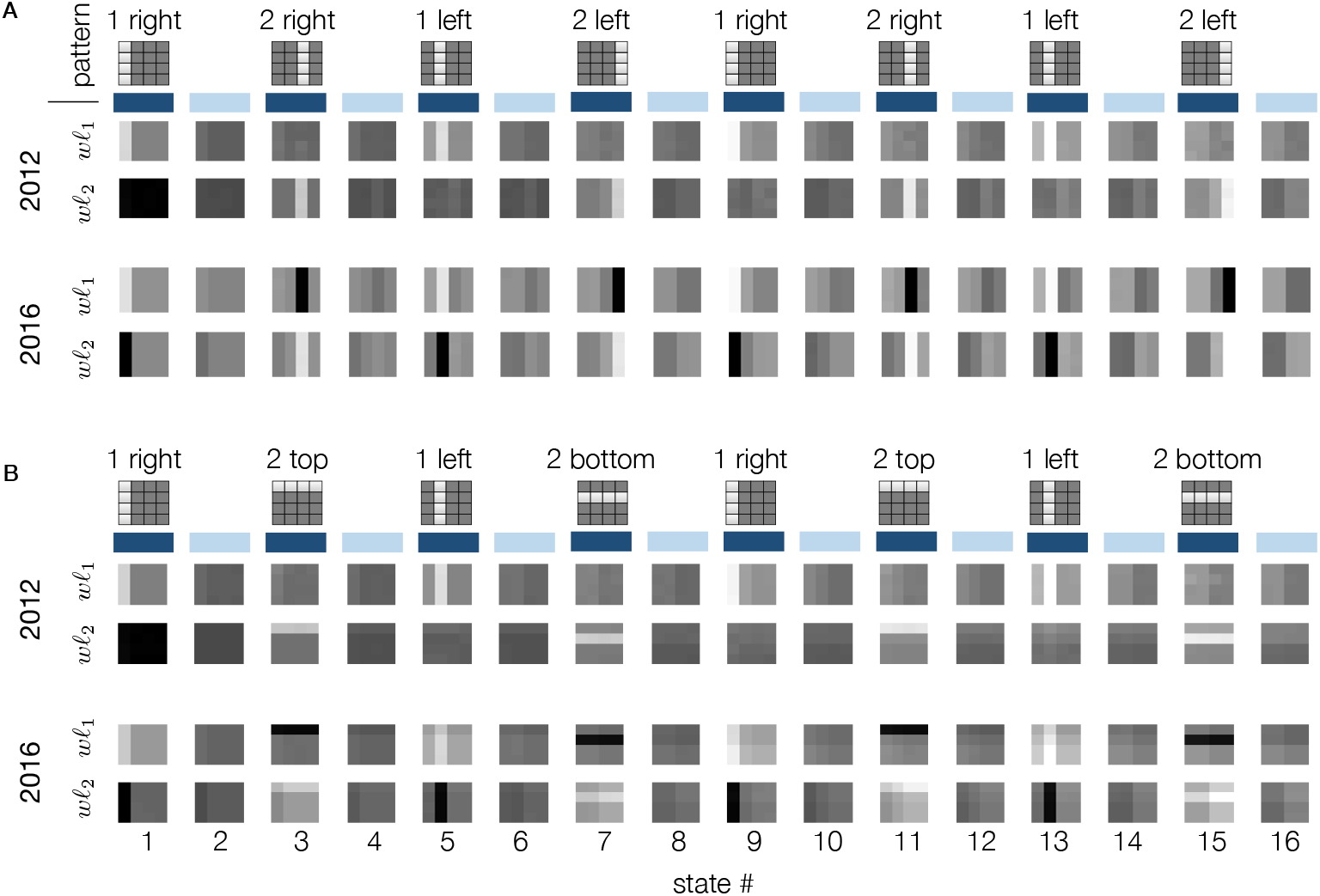
Evolution of the weight matrices for the 2 patterns at different states, associated with Figure 5, **A**. for non-overlapping patterns or **B**. overlapping patterns

