## Supplementary Material for "Two-stage synaptic plasticity enables memory consolidation during neuronal burst firing regimes"

#### Text S1 Calcium dynamics

We consider the linear calcium dynamics suggested by (Graupner and Brunel, 2012; Graupner et al., 2016). The presynaptic and postsynaptic spike contributions add linearly. Indeed, calcium enters from NMDA receptors and voltage-dependent calcium channels. Instead of describing the whole calcium, the phenomenological effect on the calcium variation is considered. At each pre or postsynaptic spike (respectively named as $t_{j,k}$ and $t_{i,k}$), the calcium immediately rises and then exponentially decays characterized by a calcium decay time constant equal to $\tau_{\text{Ca}}$:

$$\frac{dc_j}{dt} = -\frac{c_j}{\tau_{\text{Ca}}} + C_{\text{pre}} \sum_{k \in \mathcal{T}_j} \delta(t - t_{j,k} - D),$$
$$\frac{dc_i}{dt} = -\frac{c_i}{\tau_{\text{Ca}}} + C_{\text{post}} \sum_{k \in \mathcal{T}_i} \delta(t - t_{i,k}),$$

where $C_{\text{pre}}$ and $C_{\text{post}}$ are the presynaptically and postsynaptically evoked calcium amplitudes. The parameter $D$ is a time-delay between the presynaptic spike and the corresponding postsynaptic calcium transient occurrence accounts for the slow rise time of the NMDAr-mediated calcium influx (Graupner and Brunel, 2012; Graupner et al., 2016).

The total calcium amplitude $[\text{Ca}^{2+}]_{ij}(t)$ driving the synaptic change is given by:

$$[\text{Ca}^{2+}]_{ij}(t) = c_j(t) + c_i(t).$$

The time-evolution for several pre- and postsynaptic spiking activity is written such as (Graupner et al., 2016):

$$[\text{Ca}^{2+}]_{ij}(t) = \sum_{k \in \mathcal{T}_j} C_{\text{pre}} \exp\left(\frac{t - t_{j,k} - D}{\tau_{\text{Ca}}}\right) + \sum_{k \in \mathcal{T}_i} C_{\text{post}} \exp\left(\frac{t - t_{i,k}}{\tau_{\text{Ca}}}\right).$$

The resting calcium concentration is set to zero. The calcium concentrations are dimensionless. Both simplification is acknowledged because the synaptic rules are adapted in accordance. If a resting calcium concentration is wanted, the thresholds of potentiation and depression will be adapted. This notation follows the original paper notation.

#### Text S2 Computational experiments

The parameters associated with the primary synaptic plasticity are given in (Graupner and Brunel, 2012). $\tau_{\text{Ca}} = 22.6936\,\text{ms}$, $C_{\text{pre}} = 0.56$, $C_{\text{post}} = 1.24$, $D = 4.60\,\text{ms}$, $\tau_w = 346.3615 \times 10^3\,\text{ms}$, $\gamma_{\text{p}} = 725.085 \times 1.1$ (Tonic), $\gamma_{\text{p}} = 725.085 \times 0.95$ (Burst), $\gamma_{\text{d}} = 331.909$, $\theta_{\text{p}} = 1.3$, $\theta_{\text{d}} = 1$, $w^* = 0.5$. The potentiation rate $\gamma_p$ is slightly scaled up during tonic firing to induce stronger potentiation compared to the initial model, and it is reduced by 5 % during burst firing to place the reset at a lower value compared to the initial model.

The parameters used in each simulation are $N = 50, M = 50, T_{\text{state}} = 20\,\text{s}, N_{\text{state}} = 8$ for Figure 1, $N = 484, M = 10, T_{\text{state}} = 15\,\text{s}, N_{\text{state}} = 62$ for Figure 2 and Figure 3, $N = 50, M = 50, T_{\text{state}} = 20\,\text{s}, N_{\text{state}} = 4$ for Figure 4, $N = 100, M = 1, T_{\text{state}} = 20\,\text{s}, N_{\text{state}} = 10$ for Figure 4, and $N = 16, M = 2, T_{\text{state}} = 15\,\text{s}, N_{\text{state}} = 16$ for Figure 5. Some parameters remain constant in the different simulations: $I_{\text{app,inh}}(\text{Tonic}) = 3\,\text{nA/cm}^2$, $I_{\text{app,inh}}$ (Burst) $= -1.2\,\text{nA/cm}^2$, $w_0$=0.5, $\ell_0$=0.001, $\eta = 1/500$ in Figure 1, 1/400 in Figure 2, and 1/200 in Figure 5. In Figure 4, $\ell_0$ varies from 0.0001 to 0.01 and $\eta$ varies from 1/1000 to 1/10. In Figure 5, $\ell_0$ equals 0.02. To constrain the secondary weight, a maximum upper bound is present ($\ell_{\text{max}}$) - this bound is never reached in the different computational experiments. The minimum and maximum values of $w\ell$ are, respectively, $0.0265e^{-3}$, 0.00342 for Figure 2B, $0.168e^{-3}$, $0.733e^{-3}$ for Figure 2C, and $0.176e^{-3}$, $0.706e^{-3}$ for Figure 4A-B.

#### Text S3 Supplementary material related to Figure 2

Figure S1 shows the evolution of the weight matrices associated with Figure 2.

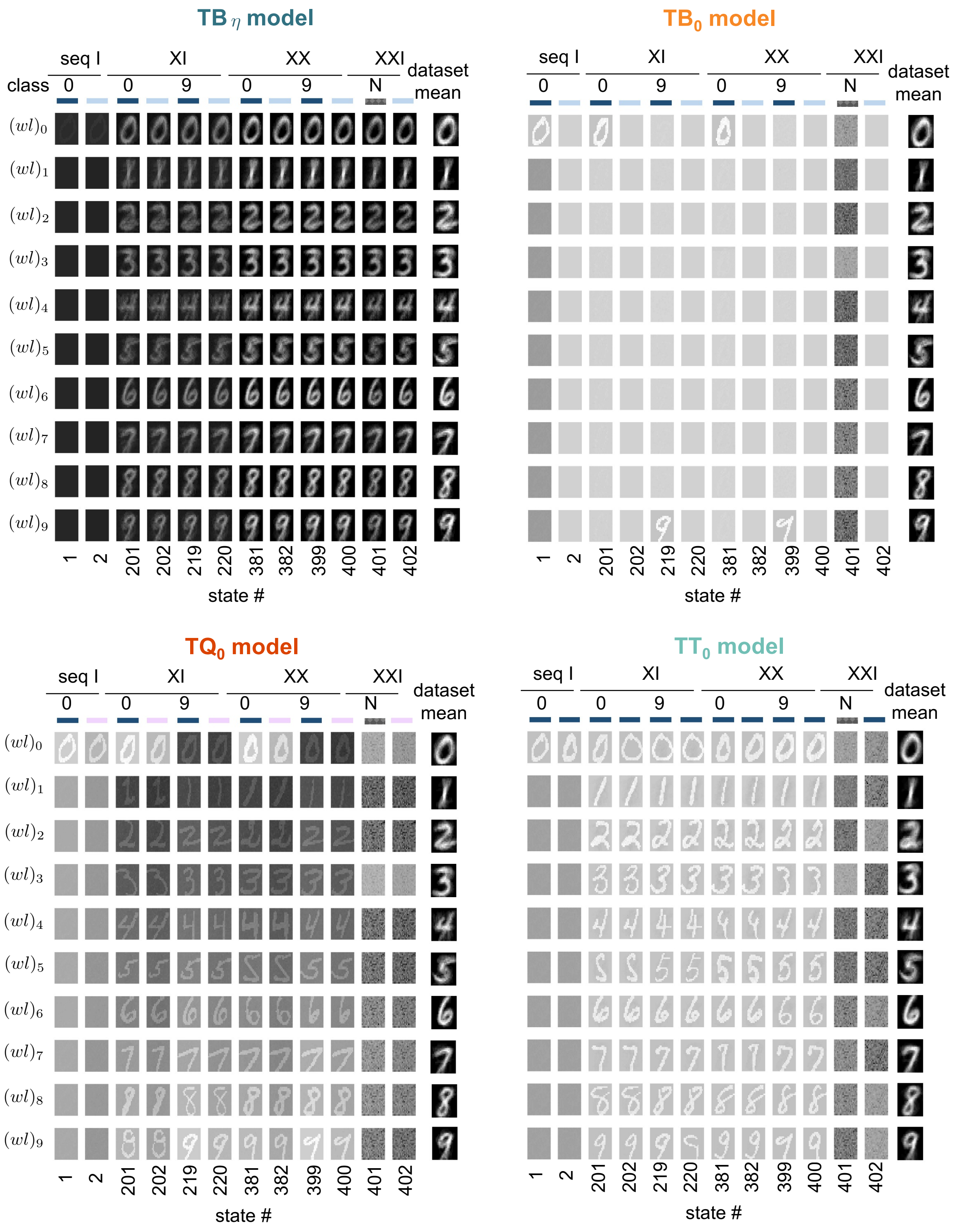


**Figure S1:** Evolution of the weight matrices for the 10 digits at different states, associated with Figure 2 for the different models (TB$_\eta$, TB$_0$, TQ$_0$,TT$_0$), depicts the protocol that interleaves different firing activities depending on the model color coded by tonic (dark blue), burst firing (light blue), noise (for N, in gray) and quiescent (lilac). (seq means sequence).

#### Text S4 Supplementary material related to Figure 3

We compare the effect of normalization on training and testing consolidation percentage. Figure S2A-B show results for unnormalized weights, while Figure S2C-D display normalized weights. Testing involves unseen samples of digits 0 to 3. Models without late-phase plasticity, where only early weights fluctuate, perform poorly, while our $TB_\eta$ model excels in the training set and can generalize its learning on the testing set. Normalization improves the performance of other models but still lags behind our approach.

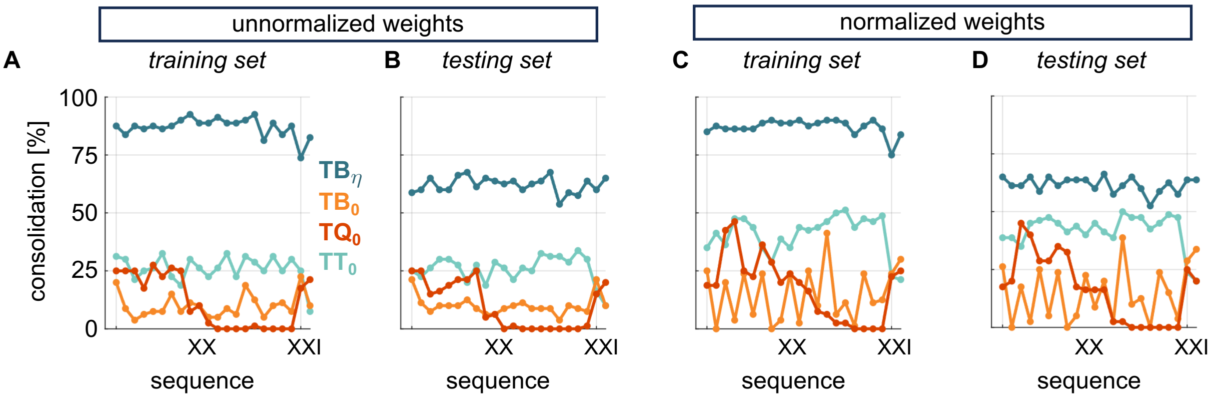


**Figure S2: Comparison of weight normalization on consolidation percentage for training and testing datasets.** Consolidation percentage was obtained for the final two sequences (XX-XXI) without weight normalization**(A, B.)**, compared to with normalized weights **(C, D)**. Consolidation percentage is compared between the training dataset **(A, C)** and the testing dataset **(B, D)**.

### Text S5 Supplementary material related to Figure 5

Figure S3 depicts the evolution of the weight matrices associated with Figure 5 across successive tonic and burst firing states. In Figure S3A, the weight matrices are compared for non-overlapping conditions between the 2012 model and the 2016 model. Similarly, Figure S3B compares the weight matrices under overlapping conditions between the 2012 model and the 2016 model.

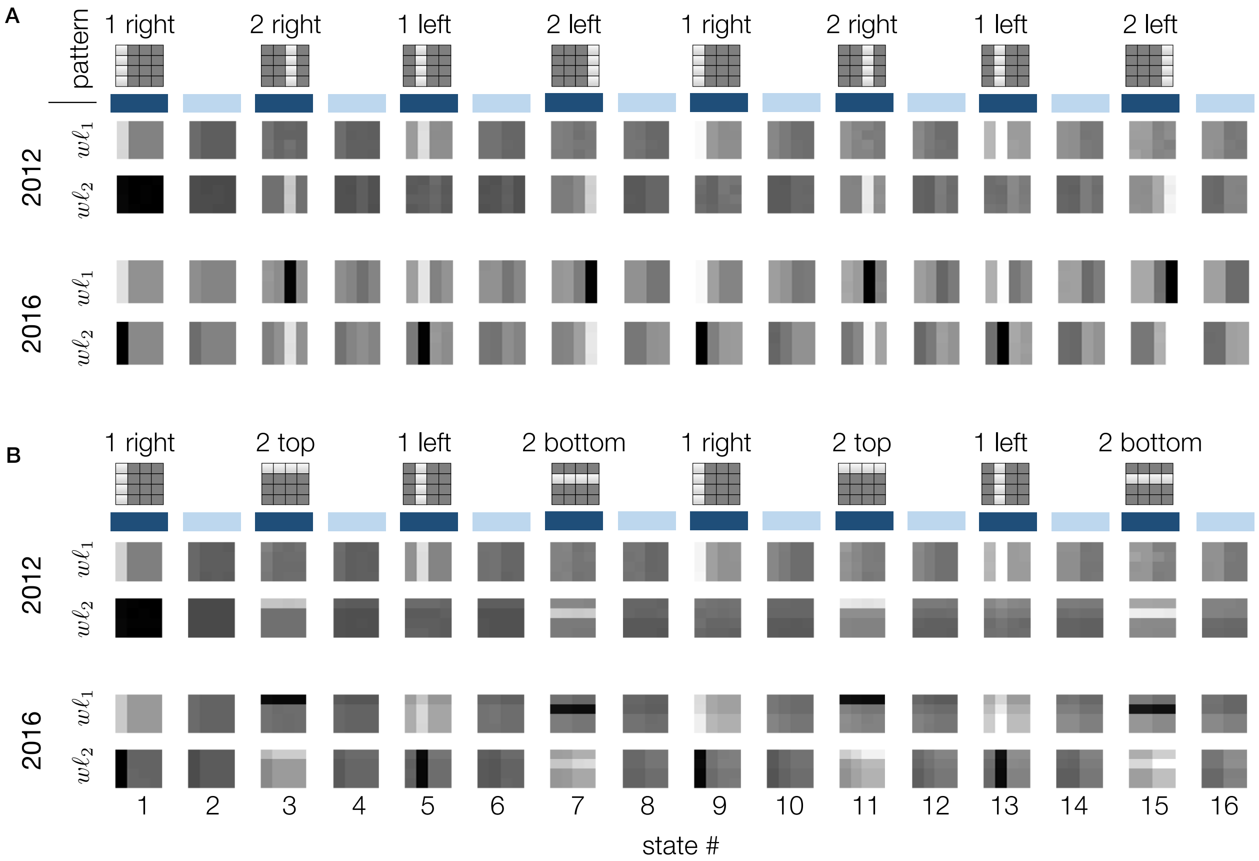


**Figure S3:** Evolution of the weight matrices for the 2 patterns at different states, associated with Figure 5, **A.** for non-overlapping patterns or **B.** overlapping patterns
